# Molecular mechanisms of GSK3*β*-driven modulation of ABLIM1 and titin interactions in cardiac muscle

**DOI:** 10.1101/2024.12.07.627363

**Authors:** Bin Sun, Alec Loftus, Brandon Beh Goh Beh, Aalaythia Hepburn, Jonathan A. Kirk, Peter M. Kekenes-Huskey

## Abstract

The heart adapts to cardiac demand through various mechanisms, including chemical modifications of myofilament proteins responsible for cell contraction. Many of these modifications, such as phosphorylation, occur in unstructured, or intrinsically disordered, regions (IDRs) of proteins. Although often challenging to study, these IDRs are increasingly recognized as dynamic, tunable regulators of protein function. Given that cardiac dysfunction can involve changes in the post-translational modification (PTM) status of myofilament proteins, it is critical to assess how alterations within these disordered regions impact intact protein and myofilament behavior. We hypothesized that the function of ABLIM1, a myofilament protein containing an important IDR, is regulated by altering its IDR conformational ensemble through PTMs, primarily phosphorylation. We proposed that this conformational change would modulate its ability to bind to other myofilament proteins. To evaluate this hypothesis, we employed a multiscale modeling approach including molecular dynamics simulations. This was used to predict the conformational ensembles of ABLIM1 before and after phosphorylation, at sites known to be altered in a canine model of heart failure with reduced GSK3*β* activity. We then used a state-based model of contraction to rationalize the physiological consequences of the molecular-scale predictions. Based on our data, we observed that local physicochemical alterations induced by phosphorylation in ABLIM1’s intrinsically disordered regions significantly affect its overall conformational ensemble properties. This ensemble change subsequently influences the ability of its LIM domains to interact with titin. Furthermore, using the contraction model, we show that a reduced ability to recruit myosin heads for cross-bridge formation, resulting from the modified LIM domain/titin interactions, provides a mechanism that elucidates previous findings of diminished length-dependent activation. These findings offer crucial molecular insights, reframing IDRs not merely as structural noise but as key, tunable elements that control protein interactions and ultimately impact mechanical behavior in the sarcomere. This work bridges molecular disorder and biomechanical function, providing a new lens to understand dynamic control and dysfunction in cardiomyocyte contraction.

## 2 Introduction

The sarcomere is the main contractile element of cardiac muscle (Fig. 1). Its tens of proteins are tightly coupled to the myofilament, where they are directly responsible for generating force in response to elevated intracellular Ca^2+^. Although the structure and function of many sarcomere proteins have been elucidated, myofilament associated proteins with intrinsic disorder (MAPIDs) [1] are among those that remain poorly understood. Much of this challenge stems from the scant availability of biophysical data for the unstructured regions. The mechanosensitive ABLIM1 protein is an example of a MAPID serving as a key mechanosensitive regulator of muscle contractility [2].

**Figure 1:**
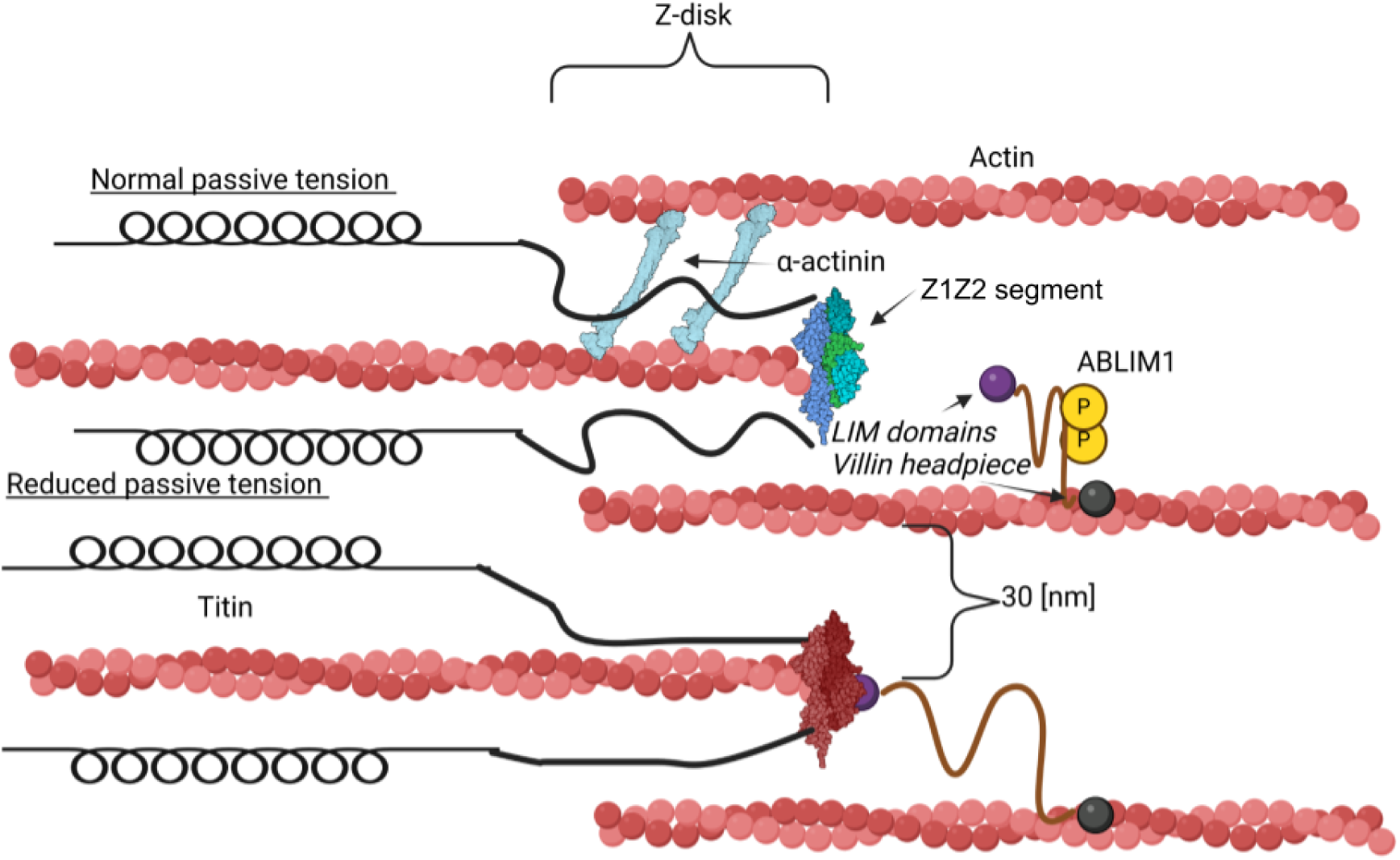
Schematic of ABLIM1 interacting with titin. This study investigates how reduced phosphorylation of ABLIM1 could affect its binding to the Z1Z2 segment of titin in the cardiac sarcomere. The C-terminus of ABLIM1 is assumed to be anchored by its villin headpiece domain (gray), while the N-terminal LIM domain region (purple sphere) can potentially interact with the titin z1-z2 domains (blue and light blue). ABLIM1 is a target for GSK3*β* phosphorylation and impacts passive tension in the myofilament [3]. The scheme was created using Biorender (www.biorender.com).

Actin-binding LIM protein 1, encoded by the ABLIM1 gene, is one of more than 60 proteins with LIM domains, which play crucial roles in the regulation of developmental pathways and cellular responses to mechanical stimuli [4]. This is done by facilitating protein-protein interactions with auxiliary proteins. Thus, the LIM domains of the ABLIM1 N-terminus have the potential to mediate interactions between actin filaments and cytoplasmic targets (Fig. 2). ABLIM1 protein (88 kD) is known to traverse the actin cytoskeleton [5]. Recent evidence suggests that it also establishes a connection between the Z-disk binding domains of titin, which is believed to contribute to the regulation of length-dependent activation (LDA) in cardiomyocytes [3].

**Figure 2:**
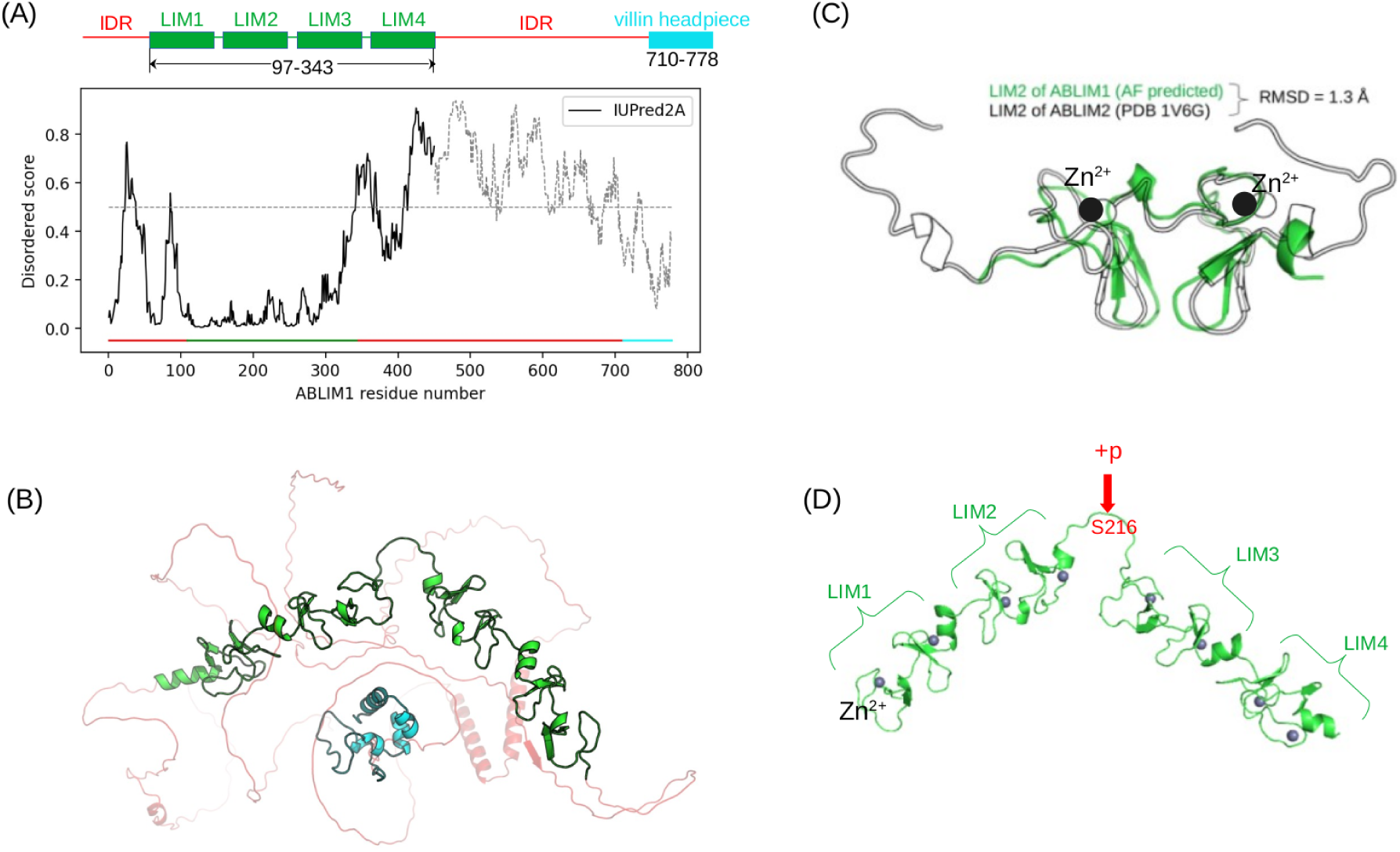
ABLIM1 Structure. A) Domain organization of human ABLIM1 protein. The disordered score of the ABLIM1 protein predicted by the IUPred2A algorithm [26]. In this study, we considered residues 1-450 of the ABLIM1 protein, and this construct consists of four tandem LIM domains flanked by two intrinsically disordered tails. B) The AlphaFold predicted structure of the entire ABLIM1 protein. C) Overlay of the LIM2 domain predicted by AF of ABLIM1 onto the experimentally resolved LIM2 domain of ABLIM2, in order to justify the use of the ABLIM1 structure predicted by AF as the starting structure for subsequent simulations. D) Zoomed in view of the four LIM domains of ABLIM1, and we consider the phosphorylation of S216 between the LIM2 and LIM3 domains.

ABLIM1 comprises four LIM domains linked to a villin headpiece by a largely intrinsically disordered region. Thus, like many other sarcomere proteins, ABLIM1 intrinsically disordered region (IDR)s are subject to post-translational modifications, which suggests their importance in the rapid regulation of contraction [1]. However, their contributions to myofilament function have not been fully resolved. An indication of its role comes from a study showing that an IDR linker in a similar LIM domain containing protein binds to plant actin to facilitate its self-aggregation, thus influencing the remodeling of F-actin [6]. Therefore, we anticipated that IDRs in ABLIM1 may play similar roles in regulating important actin-dependent functions in mammals.

ABLIM1 is known to be highly phosphorylated [5, 7], according to the PhosphoSitePlus database [8]. The C-terminal half of ABLIM1 contains several potential phosphorylation sites post-translational modification (PTM), but multiple sites are also identified in its N-terminal segment. A study [3] discovered that the state of phosphorylation of ABLIM1 affects its interactions with the Z1Z2 segment of titin, which coincides with changes in contractility. Two of these sites are located between the LIM2 and LIM3 domains (see the magenta region in Fig. 3).

**Figure 3:**
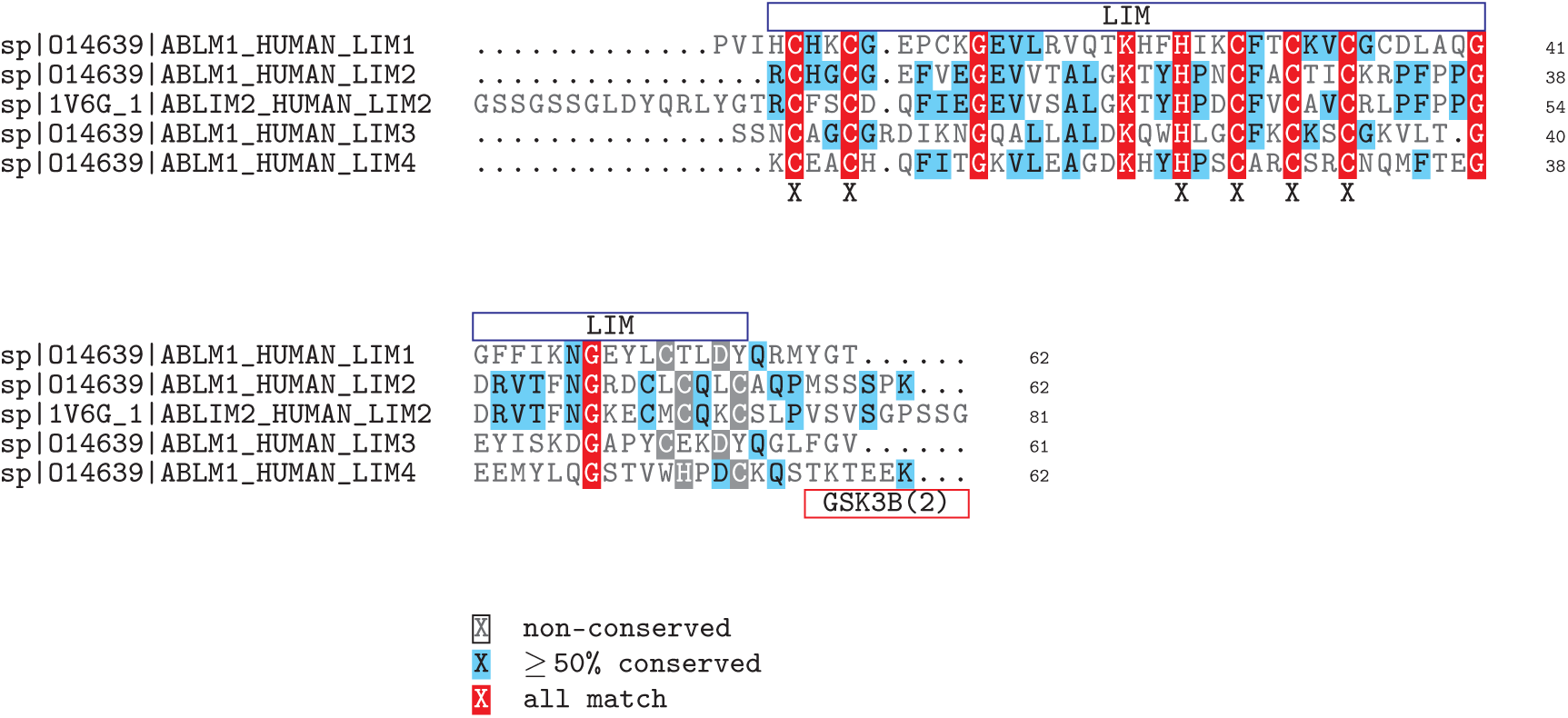
ABLIM1 and ABLIM2 LIM2 sequence alignments. A multiple sequence alignment compared the sequence structures of each ABLIM1 LIM to one another, as well as to our reference sequence, ABLIM2 LIM2. Of the red boxes where all sequences align, letters ‘C’ and ‘H’, representing Cysteine and Histidine respectively, make up the residues that bind zinc ions in the zinc-fingers of each LIM domain. Conservation between domains is summarized in Fig. S1 Generated from ClustalW [30]. An X below the alignment denotes coordination residues that are aligned in all sequences while a gray highlight denotes coordination residues not perfectly aligned.

GSK3*β* is a critical kinase that phosphorylates ABLIM1, facilitating strong force generation at extended sarcomere lengths (length-dependent activation, LDA) in healthy cardiac tissue (see Fig. 1). Its knockout in a mouse model was shown to reduce the phosphorylation of ABLIM1 at S354, S494, and S496 (see Table S1), with additional phosphorylated protein fragments remaining unchanged. A study by Stachowski *et al* [3] identified that in normal cardiac function, increased GSK3*β* activity sustains high levels of ABLIM1 phosphorylation, negligible titin/ABLIM1 interactions, and pronounced LDA. In contrast, a decrease in GSK3*β* activity typical of heart failure (HF) [3, 9] results in lower levels of phosphorylated ABLIM1, increased titin/ABLIM1 binding, and reduced LDA. Therefore, we hypothesized that the phosphorylation status of ABLIM1 impacts the ability of its LIM domains to access titin in the myofibril, where it can hinder LDA.

In this investigation, our objective was to identify how phosphorylation affects the properties of ABLIM1 in binding the Z1Z2 segment of titin. To investigate these properties, we constructed a molecular model of ABLIM1 by combining simulations of all-atom protein structures and coarse-grain representations. The all-atom simulations were performed to assess the secondary structure and key interactions that stabilize the individual LIM domains. Based on these properties, we constructed a coarse-grained model of the N-terminal domain of ABLIM1 (residues M1-S450). The coarse-grained simulations allowed us to determine the statistical properties of the ABLIM1 conformation ensemble, including the openness of the folded LIM domain regions (I97-E343) and the disordered domains (residues M1-V96 and E343-S450). Together, these simulations provide important insights into how the structure and dynamics of ABLIM1 could influence LDA, while more broadly elucidating mechanosensitive signaling of MAPIDs as a whole.

## 3 Materials and methods

### 3.1 Sequences and structures

FASTA sequences for ABLIM1 and ABLIM2 (human) were obtained from Uniprot with access IDs O14639 and Q6H8Q1, respectively. The structure derived from nuclear magnetic resonance (NMR) for ABLIM2 LIM domain 2 was obtained from the protein databank (PDB ID: 1V6G, unpublished). The structure of ABLIM2 LIM domain 2 was derived from NMR and obtained from the protein databank (PDB ID: 1V6G, unpublished). A predicted structure from AlphaFold [10] of the full-length ABLIM1 protein (human) was also downloaded from Uniprot (accession ID O14639).

### 3.2 Sequence analyses

The ELM database was used to query the ABLIM1 sequence for potential short linear interaction motifs (SLIMs), which are sequences associated with the formation of protein-protein interactions and are often subject to PTMs. The ABLIM1 FASTA sequence (from Uniprot O14639) was provided as input. The BLASTp [11] sequence alignment algorithm was used to align all LIM domains with one another to ensure sequence identity. Each LIM domain (ABLIM2 LIM2 and ABLIM1 LIMs 1-4) was treated as an independent sequence to align with all other amino acid sequences of the LIM domain.

### 3.3 Phosphorylation sites

Tandem mass spec (MS/MS) was used by Stachowski *et al* [3] to identify phosphorylated fragments from control and GSK3*β* KO myofilament-enriched myocardial tissue from mouse. In that study, phosphorylation was detected at ABLIM1 sites S215, S216, S354, S396, S397, S399, Y401, S407, S411, S466, S470, T473, S475, T477, S479, S494, T495, S496, S499, T586, S658, S670, S671, S752, and T753 (see Table S4). Only sites S354, S494, and S496 were found to be significantly different between samples (*p* ≤ 0.05), with each site having a roughly 2-fold reduction in the GSK3*β* KO tissue (Table S1).

The human sequence is 82% identical to that of the mouse, including a gap where the mouse S354 phosphorylation is found. Therefore, we modeled the equivalent positions of S450 and S452 in the human sequence (see the equivalent residues in Table S4). The protein sequences of the two species are identical in this region. Since the mouse S354 site region adjacent to LIM4 is not found in humans, we instead modeled phosphorylation of the MSSSPKET sequence found between LIM domains 2 and 3, and specifically S216, which is conserved in both species. We selected S216 because the MSSSP at site S216 (see the GSK3*β* label for ABLIM1 LIM2 in Fig. 3) is a common recognition motif for GSK3*β* and other kinases per ELM [12]. This site has been identified as a phosphorylation target in humans (PhosphoSitePlus database [8]) [13]. In addition to phosphorylation of the region identified by Stachowski *et al* [3], phosphorylation of S215/S216 has been detected in stem cells and tumors / cancers [14, 15], while a model of MLCK-null endothelial cells was found to reduce S216 phosphorylation [16].

### 3.4 Atomistic molecular dynamics simulations

We performed atomistic MD simulations of the isolated LIM2 domain of ABLIM1, the isolated LIM2 domain of ABLIM2, the LIM2-LIM3 domains of ABLIM1, the phosphomimetic LIM2 domain of ABLIM2 (S146D) and the phosphomimetic LIM2-LIM3 domains of ABLIM1 (S216D). The starting structures for these atomistic simulations were the resolved LIM2 domain structure of ABLIM2 (PDB 1V6G) or the AlphaFold-predicted full-length structure of ABLIM1. The missing Zn^2+^ cations in the AlphaFold predicted structure were added by first aligning PDB 1V6G to each individual ABLIM1 LIM domain and then copying and pasting Zn^2+^ cations into the AlphaFold predicted structure. The structure was then solvated in a TIP3P water box with a 12 Å margin and filled with 0.15 M NaCl salts. The CHARMM36m [17] force field was used to describe the system. The prepared system was first subjected to the minimization of energy for a maximum of 50,000 steps of the steepest descent algorithm (although the minimization would end early when the system reached the desired maximum force of <1.0 × 10^3^ kJ*/*mol nm^2^), followed by a heating stage during which the system was heated to 310 K over 2 ns in the NVT ensemble. During minimization and heating, restraints were applied to the protein backbone atoms with a force constant of 1 kJ*/*mol nm^2^. To maintain coordination between Zn^2+^ and coordinating residues, harmonic restraints were introduced between each Zn^2+^ and the side chains of the coordinating residues in ABLIM1. A force constant of 1 kJ*/*mol nm^2^ was applied for minimization, increased to 10 kJ*/*mol nm^2^ during the NVT equilibration step, and finally reached 100 kJ*/*mol nm^2^ for the final equilibration step and throughout production. A time step of 2 fs was used in most MD simulations, except for equilibration, where a 1 fs time step was necessary for simulation stability. The cutoff for nonbonded van der Waals interactions was set at 1.1 nm. Electrostatic interactions are handled using the reaction field method, with a dielectric constant of 15 within the cutoff distance and infinite beyond. The integration of Newtonian equations of motion used the velocity Verlet algorithm. The LINCS algorithm [18] is used for constraint calculations and the velocity-rescaling thermostat and the Parrinello-Rahman barostat are used. The velocity-rescaling thermostat has a friction coefficient of 1 ps^−1^. The Parrinello-Rahman barostat in GROMACS simulations uses a compressibility constant of 3.0 × 10^4^ 1*/*bar. The selection of Langevin over velocity-rescaling methods is driven by two factors: (1) the preservation of the canonical ensemble and (2) ensuring that the velocity distribution maintains the correct Gaussian distribution. As for the barostat, Berendsen’s equation of motion is employed to govern the evolution of the computational box’s volume and shape. Atomistic MD simulations were performed using the GROMACS2024 package [19]. We additionally performed atomistic MD simulations on the four LIM domain construct of ABLIM1, in the wild type, the phosphomimetic form (S216D), and the actual phosphorylated form (pS216), each for 500 ns production run (Sect. S1.4.2). These simulations confirm that using phosphomimetic has a similar impact as phosphorylation on the structural properties of the four LIM domain construct, both making it more extended than the WT case. More importantly, we showed that, by introducing restraints, all eight Zn^2+^ ions are well-coordinated by four atoms each, reaching the theoretical coordination number of 4 for Zn^2+^(Fig. S7), proving the effectiveness of using restraints to maintain Zn^2+^ coordination.

### 3.5 Martini coarse-grained MD simulations

To enhance sampling efficiency for large ABLIM constructs, we performed coarse-grained MD with the Martini3 [20] model. Coarse graining of the atomistic structure was conducted using the martinize.py script, and the DSSP [21] program was utilized to identify the secondary structure of the atomistic model. The CG structure was solvated with a standard Martini water model using pre-equilibrated waters from the Martini website. Elastic network constraints were applied to individual LIM domains with force constants of 500 kJ*/*mol nm^2^ and the lower and upper elastic bond cutoffs were set to 0.5 and 0.9 nm, respectively. For systems containing purely IDRs, no elastic network constraints were applied. Constraints were imposed between Zn^2+^ ions and the side chain beads of the coordinating residues of the coarse-grained (CG) models following the methodology of the atomistic simulations. The system was first energy minimized for 40,000 steps using the steepest descent algorithm and then heated to 300 K in the NVT ensemble over 1 ns. The equilibrated system was subject to a 2000 ns production MD in the NPT ensemble at 300 K with a 20 fs time step. Temperature and pressure control were maintained using the Parrinello-Rahman V scale barostat. CG simulations were conducted utilizing the Gromacs2024 package [19].

### 3.6 Continuum

A scaling relationship for intrinsically-disordered proteins was used to describe the geometric properties of IDRs within ABLIM1. This relationship takes the form of an analytic Flory random coil model [22]:

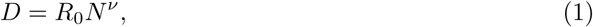

where D describes the chain dimensions, R_0_ is a prefactor in distance units, and N*^ν^* is the number of monomers raised by the apparent Flory scaling exponent. This equation has been found by others to recapitulate the geometric characteristics of proteins with IDRs [22, 23], such as for computing their effective concentrations via N^−3*ν*^[24]. This was done to determine the likelihood that ABLIM1 forms a protein-protein interaction with partners like titin, based on the protein’s ability to span two adjacent actin filaments. We used a similar approach to elucidate the binding of Troponin I to Troponin C [25]. In this paper, we compute the R*_g_* and end-to-end distance, *r_e_*, which scale as Eq. 1 according to the Flory model, in part based on fits to data from Alston et al. and Hofmann et al. [22, 23]. That is, we first determined the critical scaling factor *ν* from the values R*_g_* determined from our molecular dynamics simulations using a modified form of Eq. 1 (Eq. S1); after this, we estimated the coefficient *a*_0_ for ⟨*r_e_*⟩ = *a_o_N ^ν^*, which best fits the simulation-predicted ⟨*r_e_*⟩ for the given *ν* value. Using the ⟨*r_e_*⟩ determined from Eq. 1, we evaluated the probability of end-to-end distances using [22]:

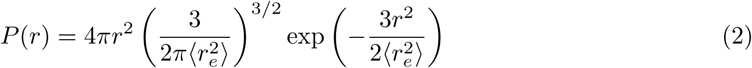

### 3.7 State modeling and fitting

Length-dependent activation was modeled using MATMyoSim [27], a software for simulating the mechanical properties of half-sarcomeres using cross-bridge distribution techniques, implemented in MATLAB [28]. These files and parameters of the tension-pCa demo were utilized to simulate the conditions of WT and mutant myocardium in Campbell et al. [29] and were adapted further to produce simulation data on trend and fitting with the observed experimental data from Stachowski-Doll et al. [3]. The parameters modified from the default tension-pCa MATMyoSim values, but consistent across all simulation models we performed, can be found in S10. The parameters that were modified for each simulation model can be found in S9.

### 3.8 Data availability

All code written in support of this publication is publicly available at https://github.com/huskeypm/pkh-lab-analyses. Simulation input files and generated data are available upon request.

## 4 Results

### 4.1 Primary sequence suggests disorder that is impacted by phosphorylation

#### 4.1.1 ABLIM1 contains globular domains

ABLIM1 comprises four LIM domains (see Fig. 2) and a single villin headpiece. Each LIM domain is stabilized by two bound Zn^2+^ ions and consists of four *β*-strands (Fig. 2C). The LIM domains exhibit 20-30% sequence identity (see Fig. 3 and Fig. S1), with highly conserved cysteine and histidine residues that coordinate Zn^2+^. Conserved glycines may confer more conformational variability to the LIM domains, given their less restrictive *ϕ* and *ψ* angles.

#### 4.1.2 ABLIM1 has intrinsic disorder

Several tools predict that many regions of the ABLIM1 sequence are intrinsically disordered. To illustrate this disorder, we show predictions from IUPred2A [26] in Fig. 2A and similar results from GlobPlot[31] in Fig. S4. This analysis reveals that almost 30% of the sequence is disordered, primarily within its N-terminus (M1-V96) as well as the segment spanning LIM4 and the villin headpiece (Q338-M696). Exceptions include folded regions within I97-E343, where its four LIM domains are located, and the C-terminus beginning with position 710, where its villin headpiece is found.

To gain insight into the topology of intrinsically disordered regions, we used the qualitative phase diagram from the Pappu laboratory [32], which relates the positive and negative fractional charge (*f_p_*/*f_n_*) to a polymer’s topology, like swollen coils. We first used PONDR-VLXT [33, 34] to predict IDR fragments in the ABLIM1 protein. Our analysis revealed several predicted IDR regions distributed in the ABLIM1 sequence, with most concentrated in the C-terminal half of the protein. This IDR distribution is consistent with the disordered scores predicted by IUPred2A as shown in Fig. 2A.

To understand how the conformational states of these IDRs would change after phosphorylation, we consulted the PhosphoSitePlus database [8] to collect all reported phosphorylation sites that fall within the predicted IDRs. As shown in Fig. 4B, the IDRs of ABLIM1 are enriched with phosphorylation sites. Upon plotting these IDRs into the Pappu diagram using their fraction of positive/negative charge fraction as coordinates, the IDRs’ projections are largely concentrated around the cusp between conformational phases, namely around the molten globules and chimeras. We then updated the *f_n_* values based on the number of phosphorylation sites we identified from PhosphoSitePlus, which shifted nearly half of the IDRs upward along the y-axis. Hence, based on the change in fractional charge, phosphorylation may shift several of the IDRs toward more extended configurations and potentially even swollen coil conformations (Fig. 4C).

**Figure 4:**
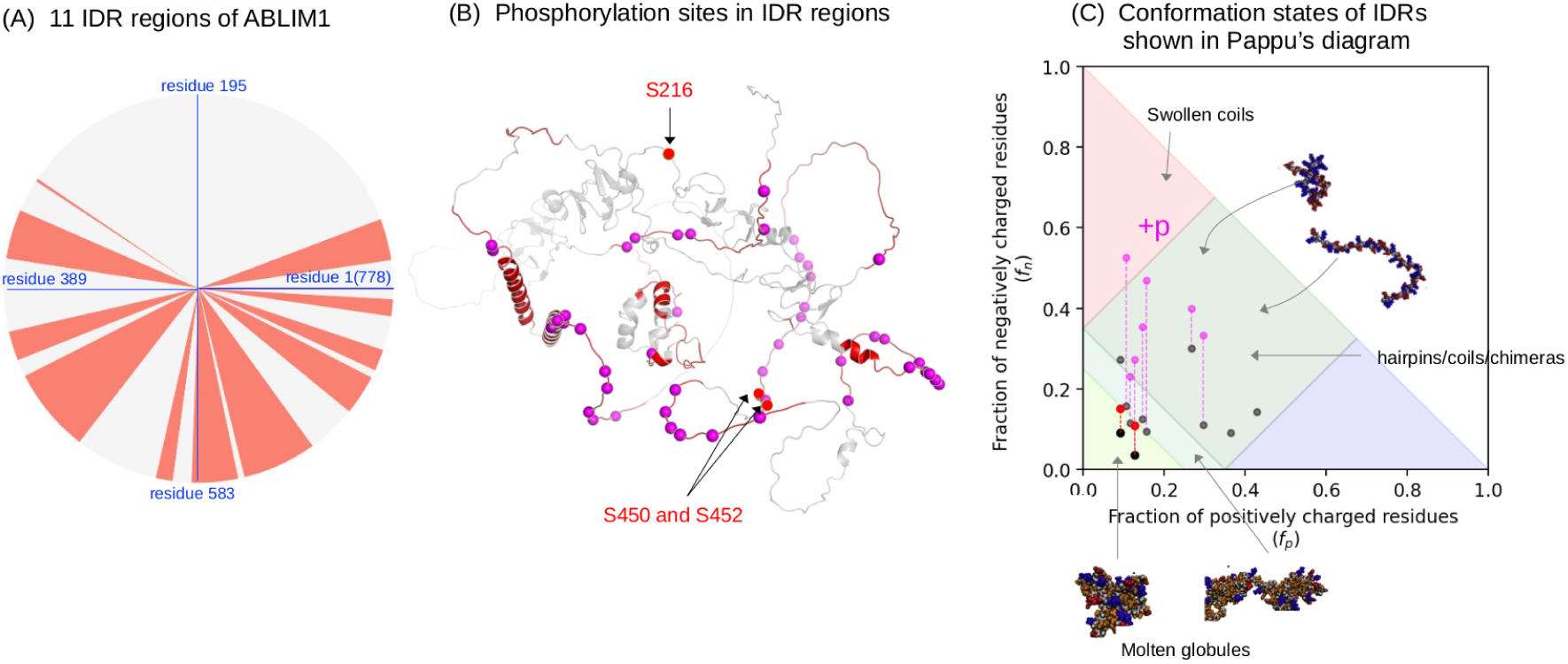
ABLIM1 Intrinsically Disordered Regions. A) Predicted IDR regions (colored red) in the ABLIM1 protein by using the PONDR-VLXT [33, 34] algorithm. The folded domains are colored white. B) Location of ABLIM1 phosphorylation sites collected from the PhosphoSitesPlus database [8], and only the phosphorylation sites located in the predicted IDRs are shown as purple spheres. The three phosphorylation sites S216, S450, and S452 that we focus on in the present study are shown as red spheres. C) Predicted topologies for native (black / yellow) IDR and phosphorylated (magenta / red) IDR based on the Pappu diagram [32]. Figures are adapted from [1].

**Figure 5:**
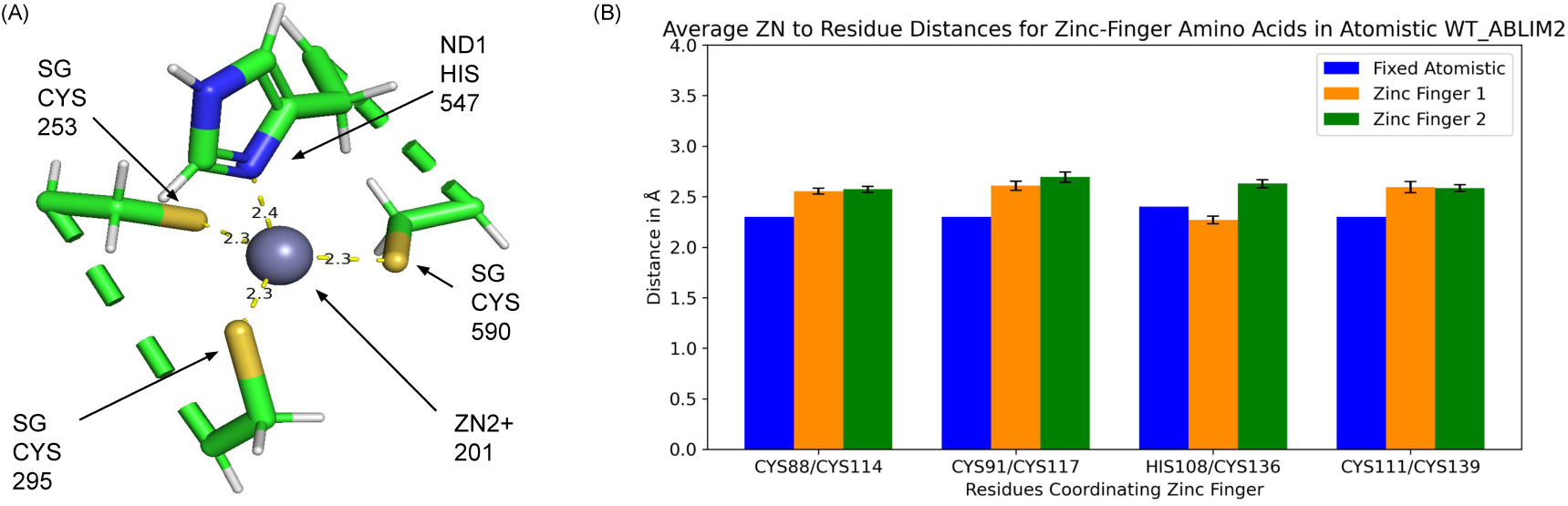
Zinc cation coordination. A) Atomistic representation of ABLIM2 LIM2 Zn^2+^ coordination structure with B) Bar chart of Zn^2+^/coordinating residue (C88, C91, H108, C111, C114, C117, C136, C139) distances computed from molecular dynamics simulations. The error bars represent the standard error of the mean distances in all atomistic simulations.

#### 4.1.3 ABLIM phosphorylation via GSK3*β* and its impact on disorder

Phosphorylation and other PTMs that alter sequence charges can shift conformation ensembles between different topologies [32]. For ABLIM, this potentially modifies its structure or interaction partners [1]. We recomputed the fractional charge density using equation *f* = *N_p,n_/N_total_*, where *N_p,n_* is the number of R/K or D/E residues, to account for phosphomimetics at three sites: S216, a putative site based on high-throughput hits [3], and two sites S450 and S452 in the disordered C-terminal region (see Fig. 4B).

These shifted sequences along the negatively charged axis (see red dots in Fig. 4C). For the three unphosphorylated sites, the IDRs containing them are located near the transition zone between molten globules and chimeras. The phosphorylation of all three sites is expected to result in the IDRs adopting more molten conformations (Fig. 4C). We anticipate that these changes will modify the orientations of the LIM domains relative to each other and will influence the effective length of the linker between the LIM domains and the villin headpiece.

### 4.2 Single LIM demonstrates little impact of phosphorylation

The sequence analyses in the previous section suggest that ABLIM1 exhibits significant intrinsic disorder, prompting us to perform molecular dynamics simulations to characterize the structure and dynamics. To begin, we utilized the atomic resolution structure available for the LIM2 domain of ABLIM2 (PDB 1V6G, unpublished structure) and the AlphaFold predicted structure for ABLIM1, as ABLIM1 does not yet have a crystallographically-determined structure available. The LIM2 domains of ABLIM1 and ABLIM2 share 67 percent sequence identity (85 percent sequence positives) according to a BLAST protein multiple sequence alignment [11] of the LIM domains (see Fig. 3). Therefore, we anticipated that the ABLIM2 structure should be a reliable surrogate for ABLIM1, given their high sequence similarity. A comparison of the structures is shown in Fig. 2. We then performed all-atom molecular dynamics simulations using ABLIM2 LIM domain 2 (Fig. 3). The binding of Zn^2+^ ions, which are important for stabilizing the LIM domains [35], was examined. For ABLIM2, Zn^2+^ ions are stabilized through cysteine, histidine, and aspartic acid residues positioned via beta sheet folds (Zinc-Finger 1: [C18, C21, H38, C41], Zinc-Finger 2: [C44, C47, C66, C69]). Consequently, we simulated the ABLIM1 fragment comprising LIM domains 2 and 3. The resulting data suggest that these interactions are maintained throughout the simulation by visual inspection for ABLIM1 as well, confirming the reliability of the atomistic model. Namely, the distances between the bonds to zinc are approximately 2.5 Å and are moderately higher than the value found for the crystal structure of LIM2 (2.3 Å).

#### 4.2.1 LIM domains are moderately labile

We report the corresponding root mean squared fluctuations (RMSF) in Fig. 6A for the ABLIM1 and ABLIM2 structures to assess the flexibility of the ABLIM structures. The smallest RMSF values are around the Zn^2+^-binding sites, consistent with the protein’s contacts with zinc ions and beta sheets remaining stable throughout the simulation. This suggests that the folded structure of the protein is relatively immobile in this region. The radius of gyration (Fig. 6 B) is relatively tightly centered about 17 Å, demonstrating a compact and well-folded structure. Calculations of R*_g_* for ABLIM1 include LIM domains 2 and 3, thus, the sizes are larger compared to the isolated LIM2 of ABLIM2. We note that the distances between the two Zn^2+^ ions assume values close to 22 Å, which aligns two of the beta sheets within the core of the protein. This configuration may be achieved through a weak hydrogen bond (approximately 4.5 Å between residues D92 and K132 seen in ABLIM2 LIM2, see Fig. 6D-E). The relative openness of this region between the two half LIM domains may indicate that the two sections could move independently of each other to bind targets. This is supported by our observation that the distribution of Zn^2+^ distance values spans 17 to 19 Å, suggesting that the protein is relatively mobile on the nanosecond timescales we simulated.

**Figure 6:**
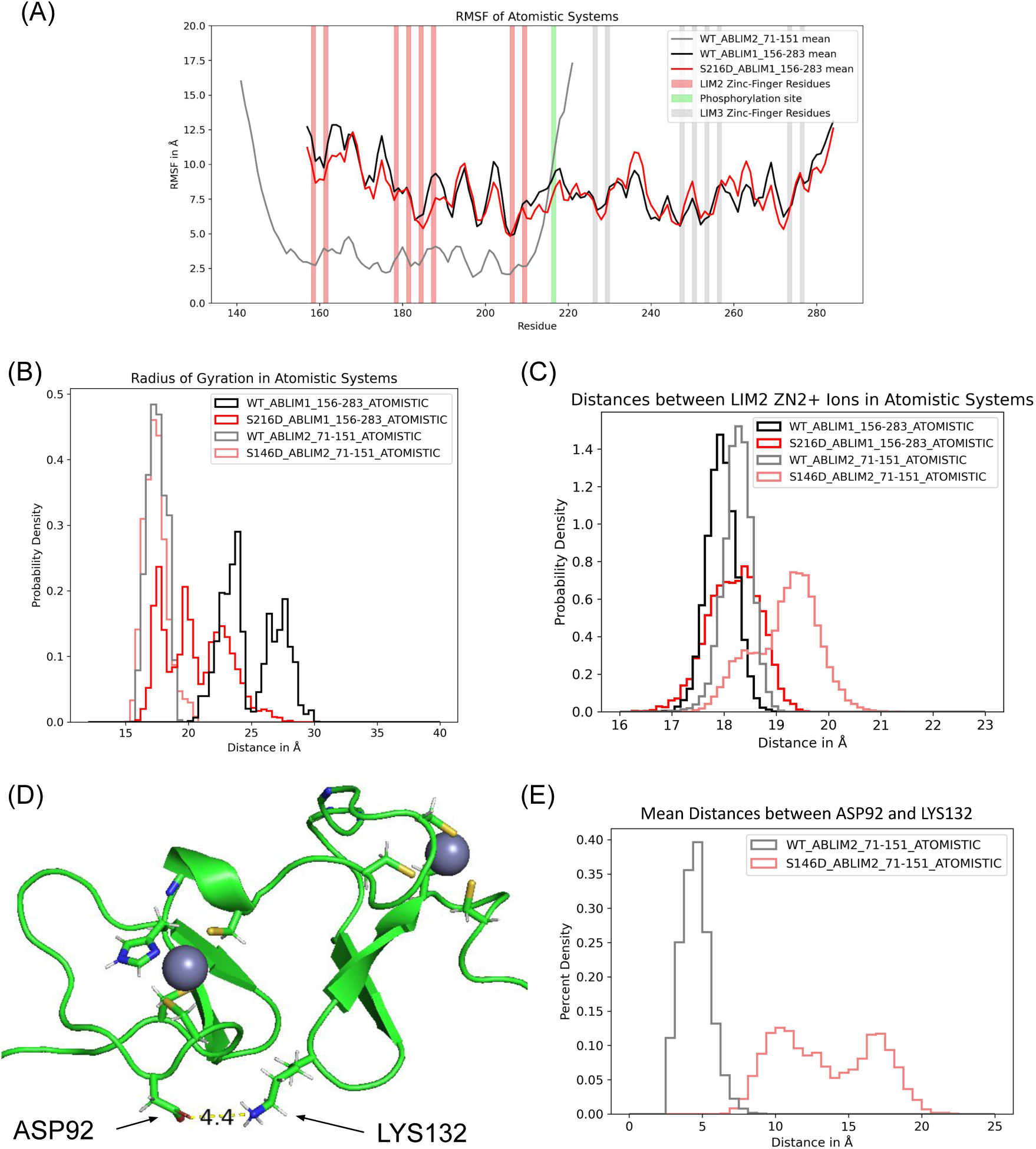
Molecular dynamics simulations of single LIM2 domain. A) RMSF comparison of atomistic systems ABLIM2 LIM2 and WT/S216D ABLIM1 LIM2-LIM3. B) Radius of gyration. C) Zn^2+^/Zn^2+^ distance showing the half LIM proximity. D) The atomistic structure of the ABLIM2 LIM domain 2, highlighting the interaction between ASP22 and LYS62 in the wild-type simulation model. E) Distance between charged elements of ASP22 and LYS62 in wild-type and S76D phosphorylated ABLIM2 simulations.

#### 4.2.2 Phosphorylation has moderate impact on ABLIM1 LIM2 structure/dynamics

We next performed simulations of the S216D phosphomimetic to approximate the structure and dynamics of ABLIM1 in its physiological, phosphorylated states relative to the (unphosphorylated) WT version. We found that the phosphomimetic at S216D did not significantly impact the RMSF trends relative to the WT (Fig. 6A), whereas the radius of gyration decreased somewhat for ABLIM1 (Fig. 6B). With respect to the distances between the Zn^2+^ atoms, phosphomimetics tended to shift them towards larger values, with smaller changes observed for the ABLIM1 construct relative to ABLIM2 (Fig. 6C). This increase in distance suggests an opening of the half LIM domains in relation to the WT.

### 4.3 LIM domain multimers are impacted by phosphorylation

In several reports [36–38], it has been observed that more than two LIM domains are generally required for LIM domain-containing proteins to bind strained actin. As an exception, testin requires a single LIM domain to bind strained actin in cells [39]. Other protein-protein interactions involving LIM domain proteins are typically mediated by their LIM domains or by additional functional domains, such as PDZ domains [4, 40]. We therefore expanded our ABLIM1 model to include all four LIM domains connected by unstructured regions, in the event that multiple domains could be used for binding to the Z1Z2 segment. Since these were unstructured, we anticipated that the LIM domains could have considerable rotational flexibility that might be insufficiently sampled by conventional atomistic simulations. Therefore, we resorted to coarse-grained (CG) Martini models to enhance conformational sampling by using larger integration time steps. Thus, we created CG Martini models of the ABLIM1 LIM domains from its atomistic equivalent (see Fig. 7A). Constraints were placed between the Zn^2+^ ions and the residues with which they coordinate; secondary structure assignments were made on the basis of results from the atomistic simulations. After equilibration and 2500 ns of simulation, the coarse-grained representations of the LIM domains remained similar to those of the atomistic data (see Fig. 7).

**Figure 7:**
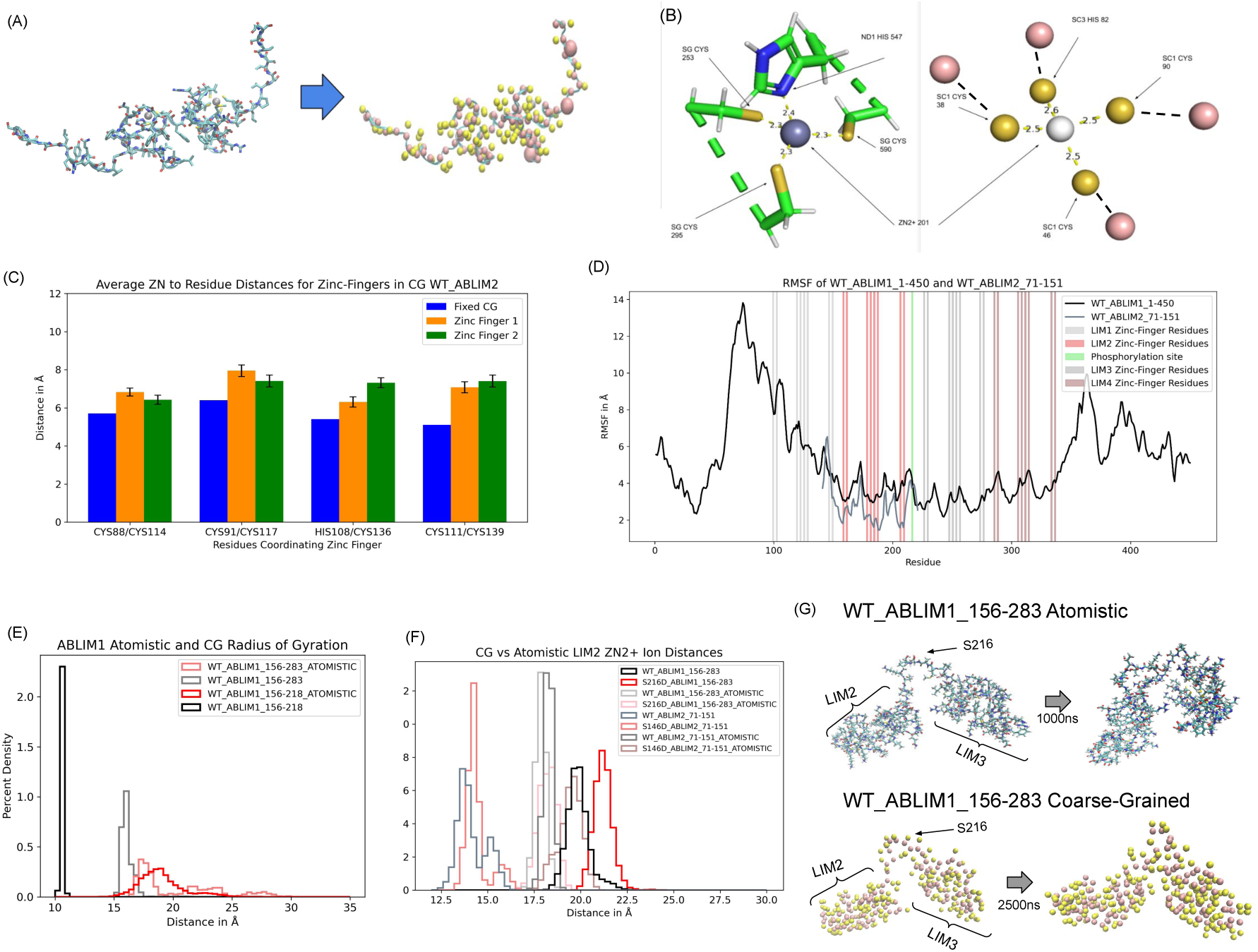
Coarse-graining. A) Mapping atomistic structure into Martini coarse-grained model. B) Zn^2+^coordination in atomistic vs. CG representations. C) Distance measurements based on B). D) RMSF of the extended coarse-grained WT ABLIM1 compared to the isolated ABLIM2 LIM2. E) R*_g_* between atomistic and CG. F) LIM2 Zn^2+^-to-Zn^2+^ distances across systems. G) Comparison of the atomic resolution and CG constructs of LIM2-LIM3 over the course of simulation.

We report in Fig. 7B the structure of the coarse-grained representation’s cation binding domain, which shows coordination similar to that in the atomistic equivalents. We report these distances in panel C, but this time using the C*α* atom of each side chain of the atomistic reference for comparison. The CG model of ABLIM maintains coordination distances similar to those in the atomistic equivalents. The beads of the CG models encompass multiple atoms, so their centers do not exactly match the side-chain distances reported in the atomistic models, which explains the greater distances in CG structures. Additionally, the CG models show a weakening of the secondary structure alignment, which slightly increases the distances between the coordinating residue side-chain beads and the zinc ions they coordinated. In addition, the RMSF shown in Fig. 7 D shows lower values near the folded LIM domains and especially near the location of the zinc finger residues, similar to the atomistic data. This suggests that our model is a faithful representation of the atomistic reference with regard to cation coordination.

Next, we examined the radius of gyration (R*_g_*) of the WT and phosphomimetic Martini models (see Fig. 7E). R*_g_* for CG are considerably smaller than the atomistic simulations, which we attribute to the different masses and positions of atomistic C*α* versus backbone beads, since the structures appear similar by visual inspection (see also panel B structures). We next describe the Zn^2+^-Zn^2+^ distances (Fig. 7 F), for which ABLIM2 LIM2 exhibited moderately smaller Zn^2+^-Zn^2+^ distances (approximately 15 Å) than WT (approximately 17Å). In comparison, their distances in LIM2 from ABLIM1 are a few Å larger than those of ABLIM2. The ABLIM1 LIM2 region (WT_ABLIM1_156-218, shown in Fig. 7F in pale blue) showed the most similar LIM2 Zn^2+^-Zn^2+^ distances to ABLIM2 LIM2, indicating that the presence of other LIM domains may impact the openness of a LIM domain. Furthermore, the phosphomimetics appear to increase this distance in all models compared to their respective WT counterparts, with the modified extended model that includes the IDRs on either end of the LIM domains (M1-V96 & E343-S450) showing the greatest increase in LIM2 Zn^2+^-Zn^2+^ distance. Despite the differences between the properties of the atomistic and CG structures, an overlay of the domains shows considerable similarity (Fig. 7).

#### 4.3.1 ABLIM1 intrinsic disorder affords diverse conformations of the 4 LIM domain construct

We extended the four LIM domain construct to include flanking regions of roughly 100 amino acids on either side. This was done to determine whether the presence of disordered flanking regions would impact the structure of ABLIM1. Consistent with the atomistic data (Fig. 6), phosphorylation did not significantly impact the values of the root mean square fluctuations (RMSF), except for a moderate reduction in the lability of the disordered N-terminus of ABLIM1 (see Fig. 8 A). The R*_g_* of the wild-type (WT) construct is approximately 25 Å, which shifted to around 28 Å upon introduction of the S216D phosphomimetic. For comparison, the WT disordered N and C-terminal regions (M1-V96 and E343-S450) assume R*_g_* of approximately 24 Å and 16 Å respectively, versus the region containing LIM domains that spans I97-E343 that was centered around 23 Å. To demonstrate the differences in R*_g_*, we provide snapshots of the WT and S216D structures in Fig. 8C, which show that the WT forms a more compact configuration. Hence, the phosphomimetic appeared to ‘straighten’ the alignment of the LIM3 and LIM4 domains, leading to distances of roughly 50 Å from S216 to site E343 for the S216D construct, versus 40 Å for the WT (Fig. 8D). We anticipated that this straightening may play a role in the binding of ABLIM1 to other target proteins. This straightening is paralleled by an increase in hydration (Fig. S2).

**Figure 8:**
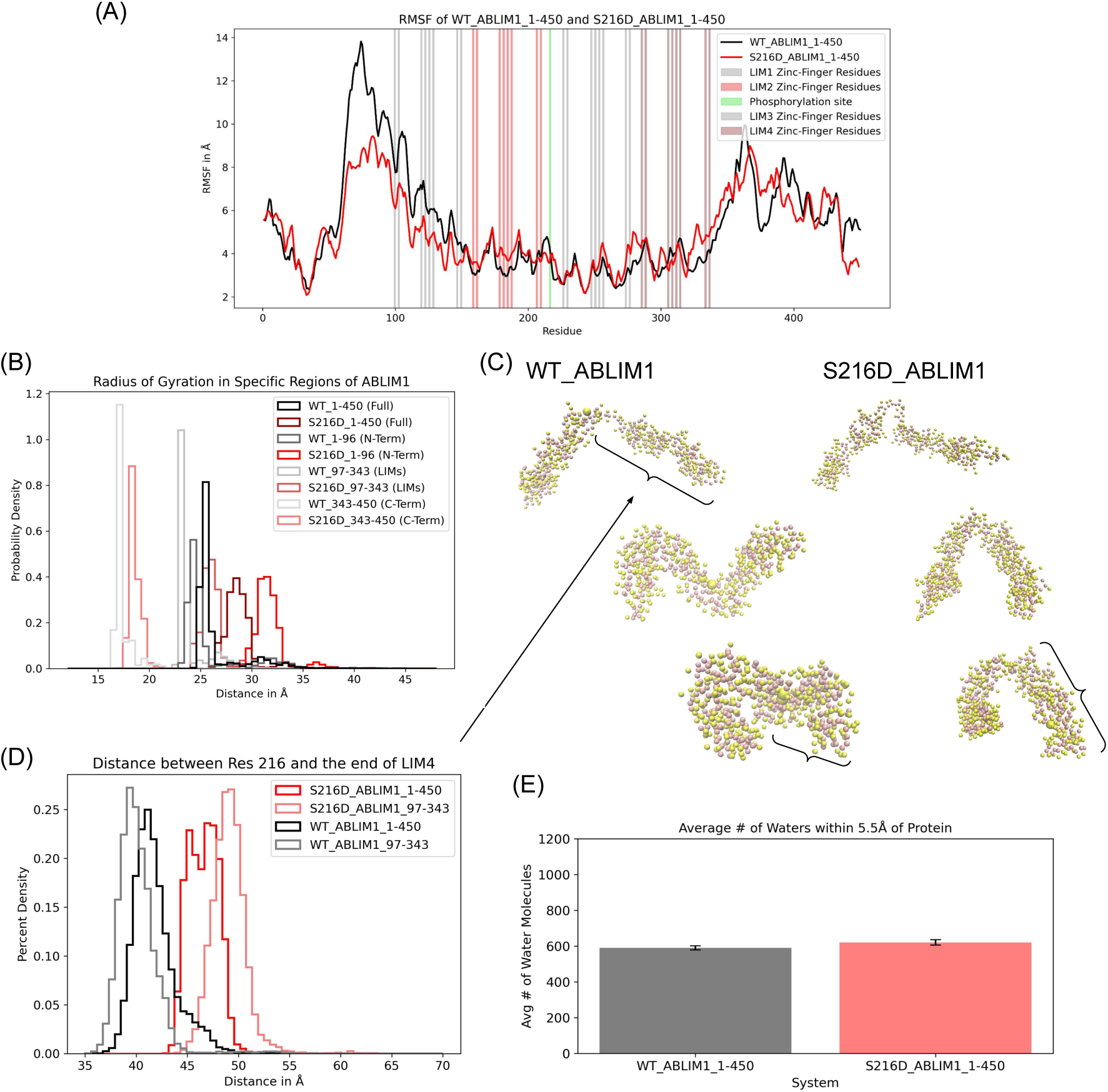
Extended CG ABLIM1 systems. A) RMSF of extended WT and phosphomimetic ABLIM1. B) radius of gyration (R*_g_*) of specific subsections of the extended simulations. C) Visualization of WT versus phosphomimetic CG ABLIM1 structures as production simulations progressed, with emphasis on the difference in LIM3-LIM4 compression from the start of the simulation to the end. D) Evaluation of compression between the phosphomimetic site and the end of LIM4 in all extended CG simulations. E) Comparison of water molecule accessibility to protein CG beads.

#### 4.3.2 ABLIM1 phosphorylation impacts its IDR conformation ensemble

We next constructed additional models with two phosphomimetics (S450D and S452D) in the disordered C-terminal region to assess whether phosphorylation consistently impacts R*_g_* for longer ABLIM1 sequences compared to the LIM1-4 construct. Simulations were then performed with the double S/T to D phosphomimetic variants. The results revealed that the double phosphomimetic impacted the radius of gyration of the isolated C-terminal disordered simulations, apparently narrowing the range of conformations assumed by ABLIM1, as shown in Fig. 9B. In addition, local differences were observed in the system, indicating that the double phosphomimetic exhibited less rigidity in the structure and allowed collapse of the structure (Fig. 9A).

**Figure 9:**
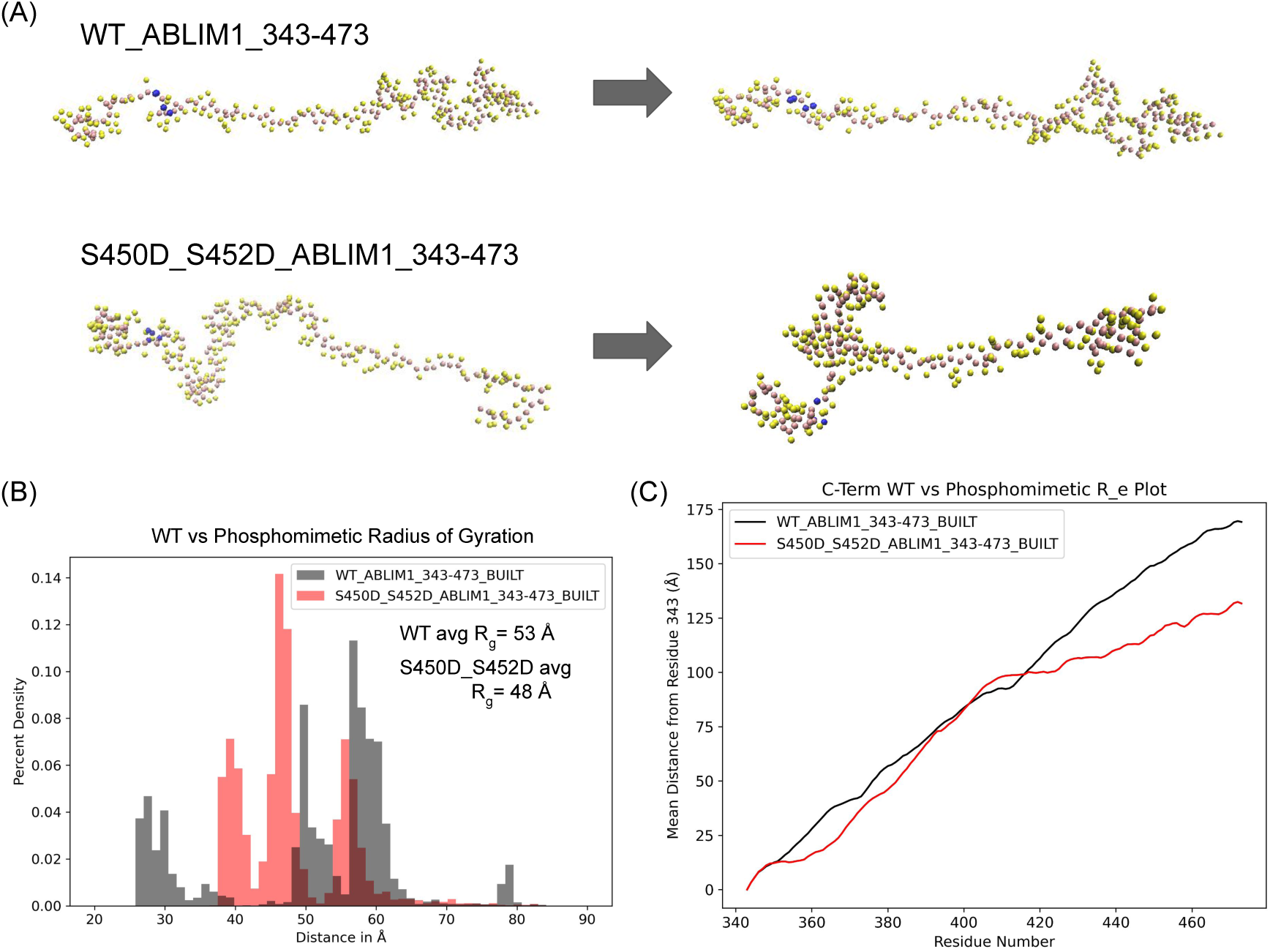
C-terminal IDR in CG ABLIM1 simulations. A) Comparisons of the WT and double phosphomimetic (S450D, S452D) structures through production simulation. B) Radius of gyration of the WT and double phosphomimetic simulations. C) End-to-end distance calculations anchored at residue E343 across both models.

Together, these data demonstrate that the WT remains extended, whereas the N-terminal region of the IDR, housing the phosphomimetics, starts to contract. Ultimately, the phosphomimetics decrease the end-to-end distance of the IDR, as shown in Fig. 9C. Evidently, the collapse of the chain near the double phosphorylation sites restricts the range of conformations that the IDR can adopt, affecting its end-to-end distance. Hence, these data suggest opposing effects of phosphorylation on different chain sizes, with increases generally observed for the LIM domain-containing region with S216D, versus a decrease observed for phosphorylation of sites S450 and S452 within the C-terminal IDR.

To confirm that our observed trends with phosphomimetics were consistent with a phosphorylated configuration, we performed simulations using explicitly phosphorylated serines (pSer). Atomistic and coarse-grained MD simulations, employing pSerine parameters adapted from Amber ff19SB [41] and Pluhackova et al. [42], respectively, both with a charge of −2e, showed that pS216 produces an extended conformation in 500 ns atomistic MD simulations on the four LIM domain construct Fig. S6, consistent with our phosphomimetic findings. Furthermore, our coarse-grained simulations that evaluate the E343-S473 fragment indicate a similar reduction in end-to-end distances of stable conformations for the protein phosphorylated at pS450/pS452 compared to the wild-type, mirroring the phosphomimetic results (Fig. S5). **rev12a** These results support the use of phosphomimetics as surrogates for constitutively phosphorylated proteins [43].

To understand the physical basis of opposing trends, we evaluated the fractional charge density of amino acid sequences near phosphorylation sites, as previously done in Fig. 4 C. For site S216, the phosphorylatable residue is flanked by the sequence motif AQPMSSSPKETTF. We selected this number of residues given that the Debye length, within which electrostatic screening effects are minimal, is approximately 1 nm at typical cellular ionic strengths [44]. The corresponding fractions of positively (*f_p_*) and negatively charged (*f_n_*) residues in the WT are 0.08 and 0.08 respectively (Table 1), which counterbalance (e.g. net charge density |*f_p_*-*f_n_*| = 0). With S216D, *f_n_* increases to 0.15, reflecting an overall increase in the net charge density. When there is a net charge due to unequal *f_p_* and *f_n_*, disordered peptides tend to have extended conformations induced by self-repulsion [45], hence explaining the straightening of the LIM domains we observed earlier (see also Fig. 4). This phenomenon is also observed for charged brush polymers that tend to swell at low ionic strength versus collapse at high salt when electrostatic screening is greatest [46].

**Table 1:**
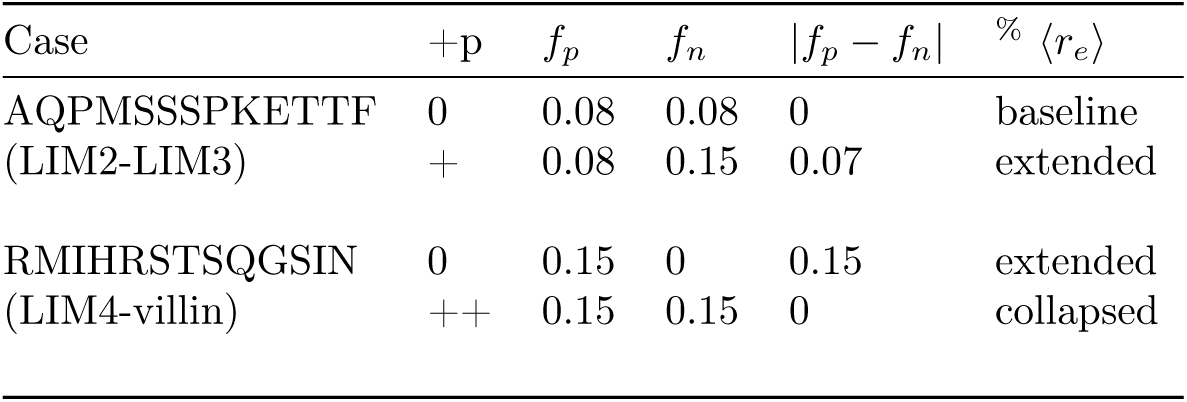
ABLIM1 fragments and their respective charge properties. The phosphorylated states are computed assuming S→D phosphomimetics. % From Fig. 11.

We applied a similar approach to examine the set of amino acids at sites S450 and S452, selecting a sequence that incorporates both residues, RMIHRSTSQGSIN. The WT form in this case had a net charge density, *f_p_* − *f_n_*, of 0.15, which is expected to exhibit self-repulsion. By mutating the two serines into aspartic acids, the charge density is neutralized. Hence, we expected that the neutralized sequence would contract somewhat relative to the WT, which is consistent with our simulation data for the C-terminal fragment (see Fig. 9C), for which the diminished electrostatic repulsion compacts the IDR. Jin *et al* [47] has also reported a similar trends Ash1, which is a positively charged 81 residue IDR with *f_p_* − *f_n_* = 0.185; namely, they reported a more compact ensemble after multisite phosphorylation. They additionally compared the effects of the IDR with a phosphomimetic (−1e) versus a protonated, phosphorylated (−1e) state and a deprotonated, phosphorylated state (− 2e); all cases rendered Ash1 more compact. Thus, the introduction of negative charges, be it from phosphorylation or phosphomimetics, to a positively charged IDR tends to compact its conformational ensemble.

### 4.4 Effects of phosphorylation of S216 on the binding of ABLIM1 to Z1Z2

Several lines of evidence suggest that ABLIM1 binds to titin [3]. Coimmunoprecipitation data from Stachowski et al. support binding in vivo, while in vitro experiments further confirmed that binding occurs between ABLIM1 and titin’s N-terminal Z1Z2 segment. Moreover, they also showed that the binding is phosphorylation-dependent, given that GSK-3*β* activity attenuates ABLIM1-Z1Z2 segment binding [3]. Based on these findings, our simulations have so far predicted that phosphorylation (S216D) opens the four LIM domain constructs relative to the WT case using CG simulations. We confirmed this observation using atomistic simulations of an ABLIM1 construct with four LIM domains (see Sect. S1.4.2).

We therefore examined the impact of phosphorylation on ABLIM1/Z1Z2 segment binding, via rigid-body docking. To this end, we selected representative conformations of ABLIM1 for both WT and S216D cases and performed ClusPro docking of these conformations to the Z1Z2 segment. This strategy was partially justified by the fact that the Z1Z2 segment is known to exist in an extended conformation in solution [48]. For each ABLIM1 conformation, we considered all poses generated by docking, of which in Fig. 10 are binding poses for the WT S1 state (the compact ABLIM1) and S216D S3 state (the extended ABLIM1 conformations). Interestingly, ABLIM1 in both cases tends to bind to the N-terminus of the Z1Z2 segment, and S216 is located far from the binding interface in all poses (Fig. S8), suggesting that phosphorylation does not likely affect binding via direct modulating the interface. We speculate that such binding poses are reasonable since the N-terminus of intact titin is a target for protein-protein interactions [49]. Among the docking poses identified, we identified several for which a *β*-strand from ABLIM1 interacts with another from Z1. The *β*-strand stacking is similar to the *β*-sheet formed at the interface in the Z1Z2 segment with telethonin (TCAP) when bound as a complex (PDB 1YA5 [49], see Fig. 12B). We further performed MMGBSA calculations on these ABLIM1-Z1Z2 segment binding poses to evaluate their stability, and we indicate that the poses exhibit favorable binding free energies (Δ*G* ≈ −40 kcal/mol, see Fig. 10), for all poses and in a phosphorylation-independent manner.

**Figure 10:**
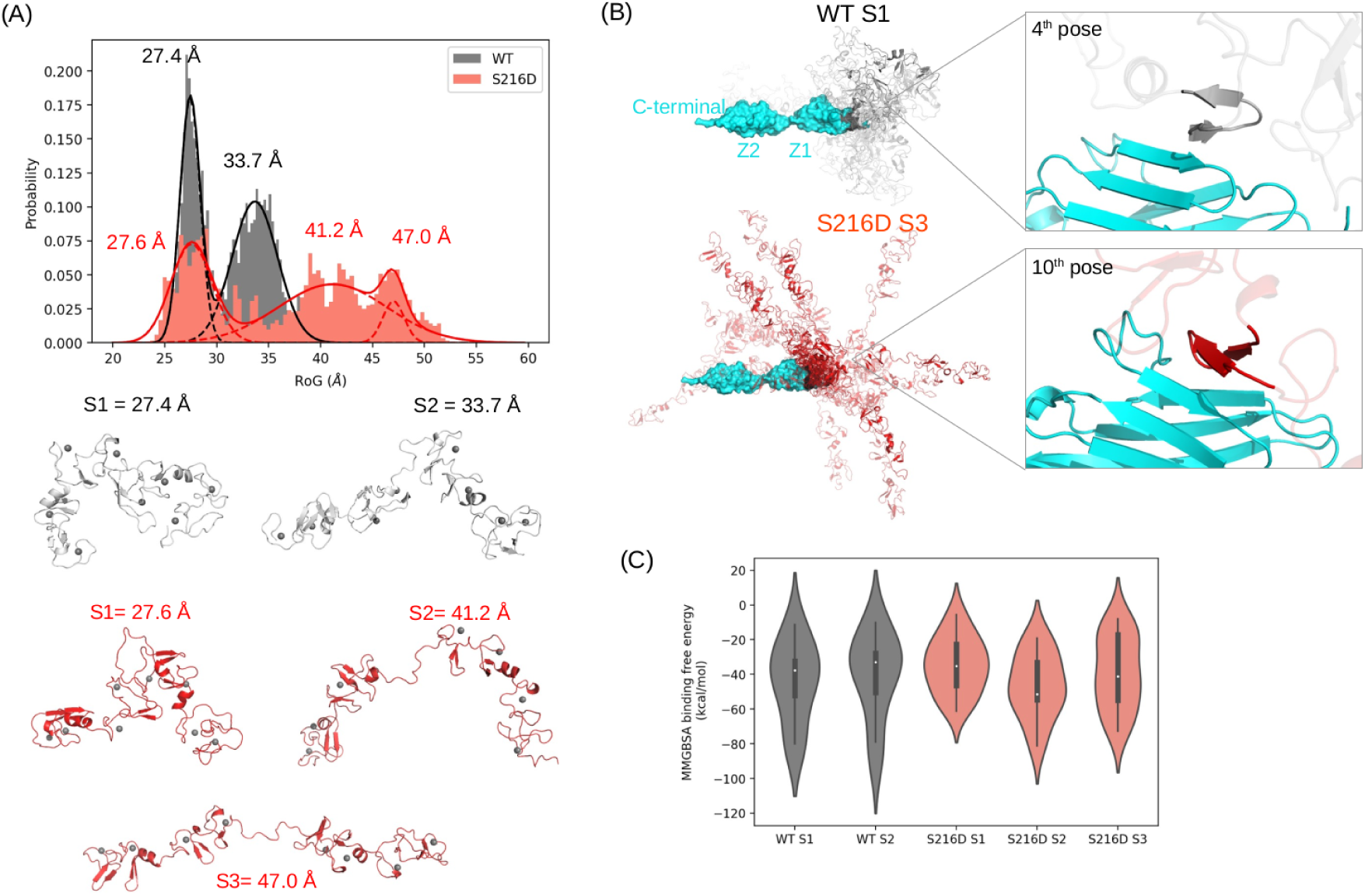
ABLIM1 Z1Z2 Docking. A) Distributions of R*_g_* of four LIM domain constructs obtained from 300 ns atomistic MD simulations (see Fig. S6). The probability density is fitted to a Gaussian function to identify the major states, and the representative conformations of the identified states are shown below. B) Rigid docking of Z1Z2 segment structure (PDB 1YA5) to the representative WT and S216D ABLIM1 structures using the ClusPro webserver [50]. ClusPro generated multiple docking poses, and we showed these poses for the WT S1 state and the S216D S3 state and also showed representative poses in which ABLIM1 forms *β*-strand interactions with the Z1 domain. C) MMGBSA-calculated binding free energy between the ABLIM1 and Z1Z2 segments based on the ClusPro generated poses.

### 4.5 Tethered ABLIM1 may interact with titin in a phosphorylation dependent manner

Our simulations and analyses of WT and phosphomimetic ABLIM1 structures suggest that the charge introduced by the phosphate groups affects the compactness of IDR. Therefore, we sought to investigate how the complete ABLIM1 could interact with titin within an intact myofibril.

To this end, we employed a Flory scaling model for polymers (Eq. 1) to predict the geometric properties of the IDR bridging LIM4 and the villin headpiece, namely the radius of gyration and the end-to-end distance. Since simulating the complete structure is not feasible, we resorted to simulations of smaller IDR ABLIM1 constructs with and without phosphomimetics to predict their radius of gyration (R*_g_*) as a function of length. This approach allowed us to estimate the scale factor *ν*, with which we can scale the radius of gyration to the full length of the ABLIM1 C-terminal IDR.

#### 4.5.1 ABLIM1 length is sufficient to overlap with titin at typical actin lattice spacings

We assumed that villin binds to a region on actin directly adjacent to the terminus of an actin from a neighboring sarcomere (see Fig. 1). This binding location is consistent with the idea that villin binds to actin [51]. The tethering of the C-terminus to actin would leave the N-terminal portion of ABLIM1 free to rotate and bind to the Z1Z2 segment dimer formed by paired titins. We next compared our sequence properties with those in Hofmann 2012 [23], where they reported data on the relationship between the radius of gyration (R*_g_*) and the number of segments (amino acids) for folded and unfolded proteins. In Fig. 11A, we compiled R*_g_* data from Hofmann *et al* [23] and our own simulations. Their study fit a form of Eq. 1 to describe this relationship for folded (orange) and unfolded proteins (blue), where the latter exhibited a larger R*_g_*. The fits for the folded and unfolded proteins were *ν* = 0.34 and *ν* = 0.58, respectively [23].

**Figure 11:**
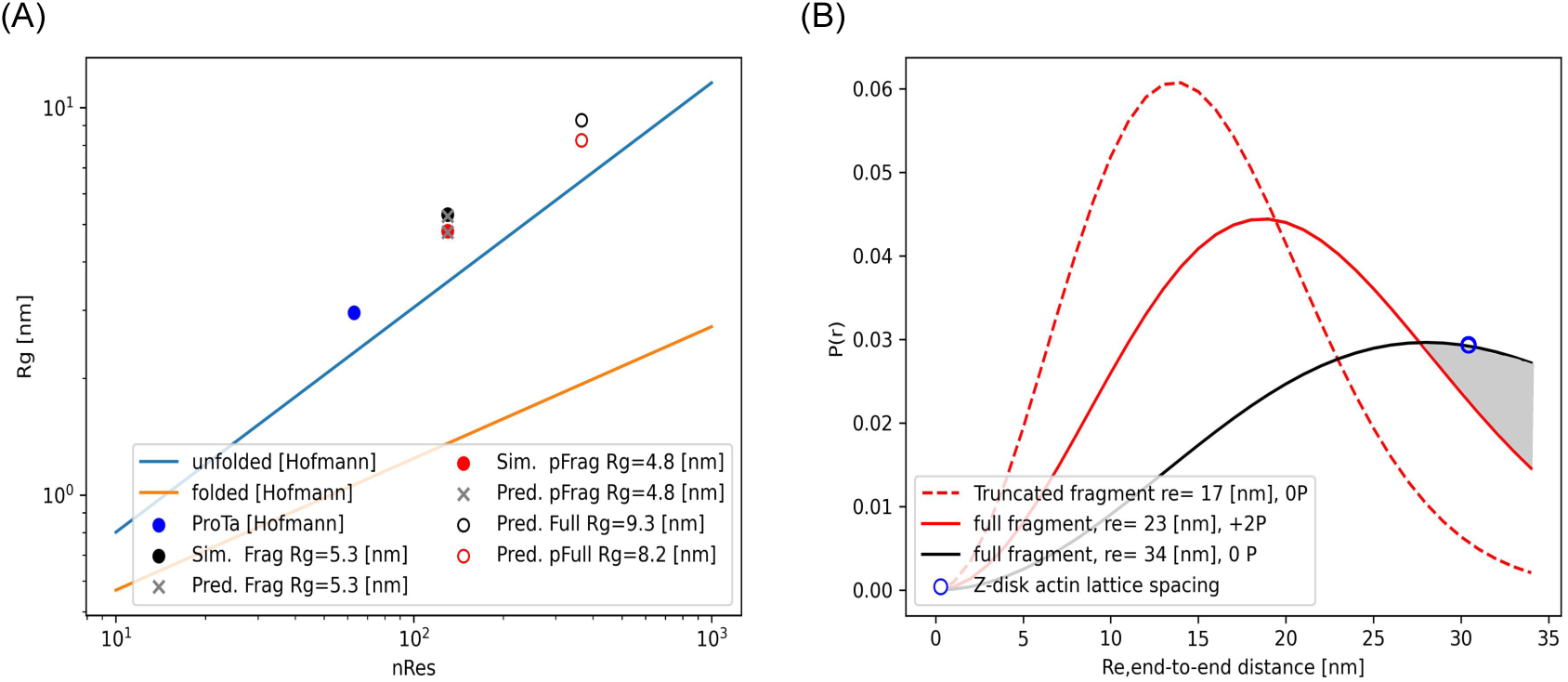
A) Radius of gyration, *R_g_*, versus number of residues (nRes), based on data from Hofmann *et al* [23] and from the molecular dynamics simulations of the truncated ABLIM1 343-473 fragment in Fig. 9. Fits to the data are extrapolated to the full-length (366 amino acid) fragment using the parameters in Table S2. B) Probability distribution of the end-to-end distances, based on the average *R_e_* estimated from panel A. An experimental estimate of the actin-actin spacing is marked in blue [52].

**Figure 12:**
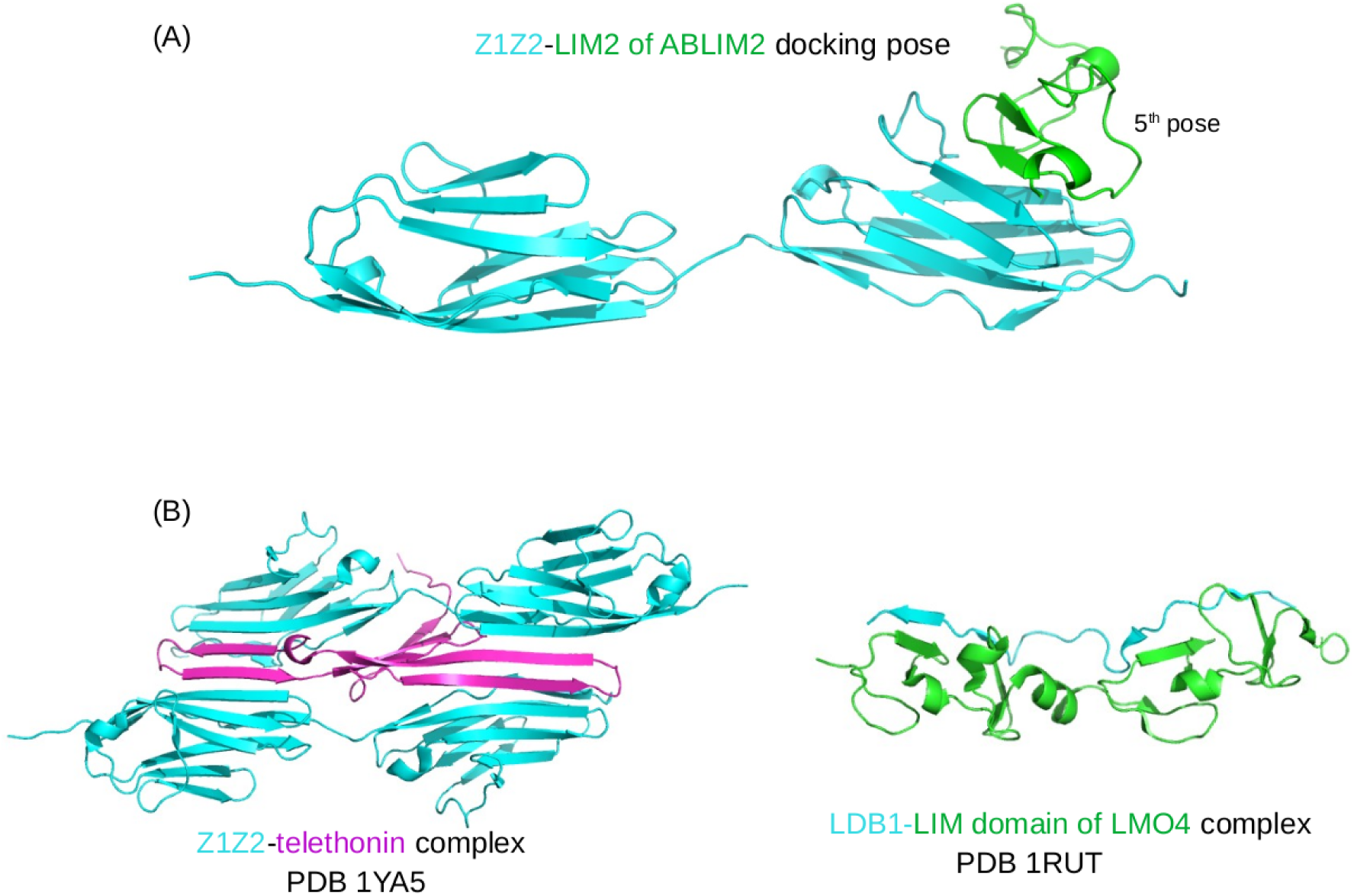
*β*-strand interaction is likely a conserved binding pattern for the LIM domain to bind to its target. A) Docking pose between the LIM2 domain of ABLIM2 with the Z1Z2 segment. B) Reported binding mode between the Z1Z2 segment and telethonin [49], and between the LIM domain protein LMO4 and its target LDB1 [36].

To compare our results with the disordered region of ABLIM1, we extracted R*_g_* estimates from our molecular dynamics simulation data. For the WT fragment spanning residues 343-473, the average radius of gyration (⟨*R_g_*⟩) of 53 Å for the 131 amino acid sequence (see Fig. 9) corresponds to *ν* = 0.68 in Eq. 1. Although this value deviates from the 36 Å estimate based on the Hofmann et al. fit for unfolded proteins, they also reported a larger than expected *R_g_* = 30 Å for a small intrinsically disordered protein ProT*α*. In that case, *ν* = 0.65 was obtained, which more closely matches our result. For our molecular simulations of the phosphomimetic fragment that yielded a ⟨*R_g_*⟩ of 48 Å, we obtained a moderately smaller scale factor of *ν* = 0.65. This decrease is attributed to the two S to D mutations neutralizing the positive charge density of the WT, mitigating swelling due to self-repulsion. Using our estimates for *ν* based on the simulated fragments, we predicted the corresponding ⟨*r_e_*⟩ values for the full (366 amino acid) C-terminal sequence as 93 and 82 Å for WT and phosphomimetic forms, respectively.

To relate the geometries of the ABLIM1 fragments to their ability to interact with the Z1Z2 segment on neighboring actin filaments, we use the probability distribution *P* (*r*) from Eq. 2. This relationship determines the likelihood that a fragment assumes a distance comparable to the spacing of the actin filaments. We first compute ⟨*r_e_*⟩ for the intact C-terminus of ABLIM1 using predictions from our molecular simulations of shorter fragments. For this, we report in Fig. S3 the ⟨*r_e_*⟩ values for the short WT and phosphorylated ABLIM1 C-terminal fragments, which were found to be 170 Å and 132 Å respectively from Fig. 9C. Using Eq. 1 again, but this time fixing *ν* to the values determined from the R*_g_* fits, we solved for *R*_0_ using the ⟨*r_e_*⟩ values from our simulation data. Our fit with the fixed *ν* values yielded *R*_0_ = 6.0 Å for linear scaling of the end-to-end distance versus the number of residues in Fig. S3. The fitted parameters provided reasonable estimates of the ⟨*r_e_*⟩ for our simulated fragments, though they under- and overestimated the values to an acceptable degree. Extrapolated fits to the Flory equation Eq. 1 for the 366 amino acid C-terminus resulted in ⟨*r_e_*⟩ values of 311 and 276 Å, respectively.

We then evaluated how the average ⟨*r_e_*⟩ influenced the chain distributions of the wild-type (WT) fragment and its phosphomimetic counterparts. These analyses yielded probability distributions for each end-to-end distance, as shown in Fig. 11. To obtain these results, we substituted the estimated average end-to-end distance (⟨*r_e_*⟩=170 Å) for the WT fragment into equation Eq. 2, yielding a maximum at approximately *r_e_*=120 Å. Based on ⟨*r_e_*⟩ values of 340 and 230 Å for the full-length C-terminal cases, we observed that *P* (*r*) maxima was right-shifted toward larger lengths of roughly *R_e_* =190 Å for the phosphomimetic case versus about 270 Å for the WT.

Lastly, we investigated the properties of the ABLIM1 protein when bound by its villin domain to an actin filament, relative to the spacing between the actin filaments (see the schematic in Fig. 1). Namely, we estimated the end-to-end distance from ABLIM’s bound villin domain to its LIM domains to assess the LIM domains’ likelihood of interacting with an adjacent actin filament capped with titin Z1Z2 segment segments. The actin lattice spacing in the Z band was estimated to be approximately 30.4 nm (Irving et al [52]), which is marked in Fig. 11B by a blue circle. The *P* (*r*) data in Fig. 11B suggest a marginally higher probability for the wild-type (WT) C-terminal region versus the phosphorylated condition will assume a value *R_e_* that matches the filament spacing. At this distance, the WT ABLIM1 is more likely to interact with a Z1Z2 segment dimer on an adjacent actin relative to its phosphorylated counterpart; by extension, the WT (unphosphorylated) case may have a greater potential to interfere with the Z1Z2 segments that anchor titin to actin. Next, we assumed that ABLIM1 is no longer perpendicular to actin but rather anchored at an angle similar to *α*-actinin (see Fig. 1); in this scenario, larger *R_e_* values would be needed to bridge adjacent filaments, which results in the WT being increasingly favored over the phosphorylated ABLIM, as illustrated by the shaded region in Fig. 11B.

### 4.6 Length-Dependent Activation (LDA)

Data from Stachowski *et al* [3] suggested that suppressing ABLIM1 expression increases force, ABLIM1 phosphorylation decreases force, and reduced ABLIM1 phosphorylation diminishes length-dependent activation. We assume that the bases of these effects stem from ABLIM1’s impacts on passive tension, namely from disrupting its interactions with the Z1Z2 segment. To link these effects to LDA, we used MyoSim [29] to simulate the maximum force generated within a cardiac sarcomere at fixed lengths and calcium (Ca^2+^) concentrations. In this model, the recruitment of myosin from its bound state on the thick filament to an off-state ready for cross-bridge formation is given by

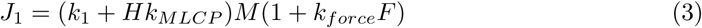

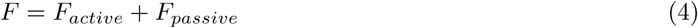

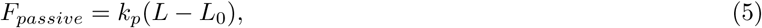

where *k*_1_ is a rate constant, *H* = 1 as the myosin regulatory light chains (RLCs) are all phosphorylated, *k_MLCP_* and *k_force_* are constants, *M* is the proportion of myosin heads in the off state, *F* is the total of the *F_active_* and *F_passive_* forces, where *k_p_* is the stiffness of the passive elastic element, *L* is the sarcomere length, and *L*_0_ is the sarcomere length at which the passive force is zero (see Fig. 13A). Using this model and parameters in S9 and S10, we varied the sarcomere lengths from 1.9 *µ*m to 2.3 *µ*m over Ca^2+^ concentrations ranging from below 0.100 *µ*M to 100 *µ*M to estimate steady-state isometric forces under resting and saturated conditions, respectively. In Fig. 13B we present the simulated contractile force as a stress (F/A), with each normalized to their maximum force at saturating Ca^2+^. We report that the predictions for the stretched sarcomere of length 2.3 *µ*m presented larger stresses compared to the resting length 1.9 *µ*m (Fig. 13B). Using the definition of the pCa_50_ as the Ca^2+^ concentration at which half-maximal force is obtained, we report values of 5.66 and 5.92 for the slack and stretched configurations, which corresponds to a left-shift of ΔpCa_50_ by 0.26 units. This shift to the left represents an increase in apparent calcium affinity, which together with the increased maximal stress observed at long sarcomere lengths affirms the Frank-Starling effect. In effect, fewer calcium equivalents were required by the stretched sarcomere to achieve similar levels of stress generated for the resting sarcomere.

**Figure 13:**
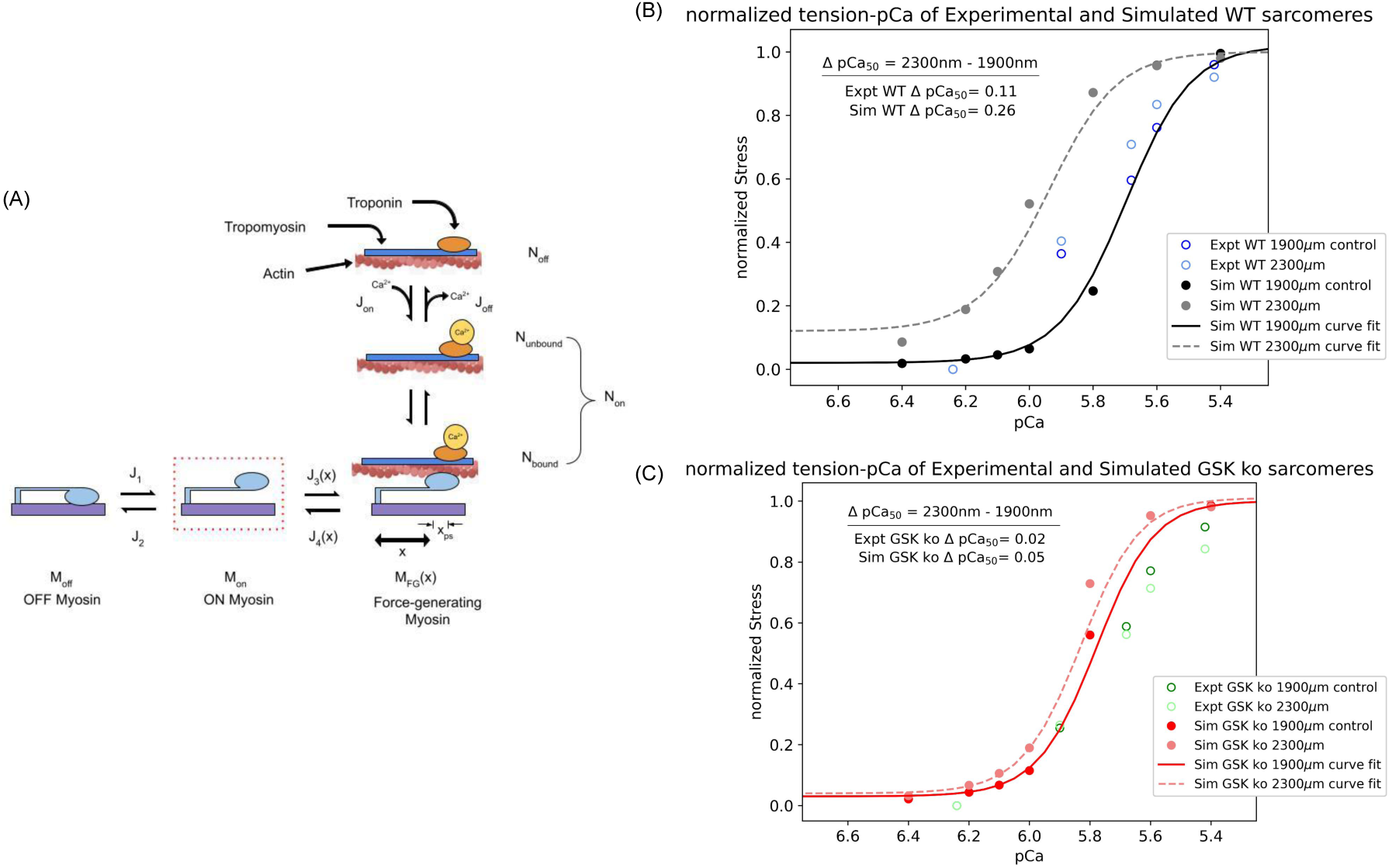
Force generation in a cardiac sarcomere. A) A kinetic model of force generation in myofibrils that assumes off, on, and cross-bridge generating states for myosin [29]. We assume that changes in passive force due to altered titin-actin interactions impact the transition rate between myosin off and on states. B) Predictions of stress (force/area) for isometric contraction at two sarcomere lengths, based on the model in panel A.

To reflect disrupted passive stiffness attributed to titin similar to the reduced activity of GSK3*β*, we reduced *k_p_* and *L*_0_ for our GSK3*β* knockout model configuration to match the experimental trends reported by Stachowski *et al* (Table S9). For this configuration, we observed a comparable maximum stress for the resting sarcomere lengths, consistent with data from Stachowski *et al*, when coupled with a change in the *k*_1_ rate constant from 1 to 2, although our results show a loss of cooperativity evidenced by the smaller slope of the stress/pCa curve. The similar level of stress despite a weaker passive force term is due to our assumption that the resting sarcomere length was shortened. We based our assumption on a similar treatment for titin mutants in [29]. We found that simulations of the GSK3*β* knockout predicted a smaller increase in maximal force at the extended sarcomere length compared to WT. Consequently, the values of Δ pCa50 were reduced by two thirds to 0.05. In other words, reducing the passive tension in our model was sufficient to reflect the loss of LDA reported by Stachowski *et al*, despite some quantitative differences in cooperativity, which could be attributed to the simplicity of the kinetic scheme used.

We note that the Campbell model [29] exhibits elevated force at low pCa (<6.2) for stretched tissue, which is consistent with force/pCa data collected by Patel *et al* [53] using rat skinned trabeculae. However, the data collected by Stachowski *et al* used isolated skinned mouse cardiac myocytes, for which the forces at low pCa converged to nearly zero at low pCa. Because we limited our changes to the Campbell model to the parameters governing the myosin off-to-on transition and slack length, our predicted tension for stretched sarcomeres in trabeculae at low Ca^2+^ is elevated relative to the slack condition, in contrast to the data from Stachowski *et al* for isolated myocytes. We did not further refine the model given the many differences in passive tension and force development in intact tissue compared to isolated cells that would be difficult to reformulate. Nonetheless, both preparations exhibit loss of LDA when the ability to generate passive force is reduced relative to WT, thus validating our hypothesis.

## 5 Discussion

### 5.1 Main highlights

In this computational study, we examined the myofilament protein ABLIM1 and its interactions with titin at the Z1Z2 segment. Our findings suggest that ABLIM1 LIM domains are compatible with binding titin, and that phosphorylation can influence the conformation distribution of ABLIM1 relative to the WT (unmodified) protein. Furthermore, intact ABLIM1 is capable of spanning typical actin-actin spacings at the z-disk, and its span is sensitive to the phosphorylation state. These molecular findings provide a mechanism linking phosphorylation-dependent ABLIM1 conformation and titin interaction to the modulation of passive tension and, consequently, length-dependent activation (LDA). Overall, these data have significant implications for mechanosignal transduction in cardiac myocytes, particularly in the context of failing hearts where reduced ABLIM1 phosphorylation has been observed [3]. These results also strengthen the support for LIM domains in mediating diverse mechanosensitive functions [2].

### 5.2 ABLIM1 binding partners

We elucidated the binding properties of ABLIM1 by analyzing the STRING database. Our search of the database revealed that ABLIM1 binds to other sarcomere proteins, including titin, *β*-actin, and LIM-nebulette, with varying degrees of confidence [54]. This finding is consistent with previous research that indicates that ABLIM proteins are generally associated with binding actin, often through their villin-type "headpiece" domain common to tens of cytoskeletal proteins [55]. Additionally, LIM domains, such as those present in ABLIM1, have been suggested to act as protein-binding modules [56] and may recognize different actin conformations, particularly when actin is stressed. Recent molecular dynamics simulations have provided a mechanism by which LIM domains can bind strained actin conformations, including the recognition of two binding sites exposed at the cracked interface [57].

Intriguingly, we identified potential associations of ABLIM1 with KCNJ12 and CLCN1 by STRING analysis. KCNJ12 codes for an ATP-dependent inward rectifying potassium channel (Kir2.2), and CLCN1 is a gene for a chloride channel. ABLIM1 has previously been identified as an actin-binding LIM protein associated with Kir2.2 in brain proteins [58]. Therefore, the expression of Kir2.2 in the heart [59] could exhibit similar interactions. A previous study indicated that CLCN1 and ABLIM1 are spliced by the same protein MBNL1 [60], but evidence for coexpression and association is lacking. The potential association between ABLIM1 and these ion channels may imply a role for ABLIM1 in modulating cell excitability. This could provide a link between mechanics and excitation at the z-disk, although more research is needed to confirm this hypothesis.

The binding of titin through its N-terminal Z1Z2 segments has been observed [3], although the specific region responsible for this interaction remains unknown. Our data suggest that LIM domains have the capacity to bind the Z1Z2 segment directly, as individual LIM domains can bind to the Z1Z2 segment of titin, as shown via our docking simulations and MMGBSA calculations. In fact, some of the predicted binding poses align a LIM domain *β*-strand with those of the Z1Z2 segment, similar to the resolved Z1Z2-telethonin complex (PDB 1YA5) where intermolecular *β*-strands interact tightly as antiparallel sheets [49] (Fig. 12). This binding mode is also reported for LMO4 and LDB1 (1RUT) [36], where *β* strands of the LIM domains align antiparallel with those of the target protein. The telethonin-Z1Z2 segment complex suggests that telethonin dimerizes Z1Z2 segments from two titin strands, bridging the *β*-sheets between a Z1Z2 pair (Fig. 12). In this capacity, ABLIM1 could compete with or compromise the binding of telethonin to the Z1Z2 segment. Consequently, telethonin mutations have already been established in hypertrophic and dilated cardiomyopathies [61]. This raises the possibility that the disruption of the binding of TCAP to titin by ABLIM1 could contribute to cardiac dysfunction.

We show that phosphorylation of ABLIM1 may play a crucial role in determining its conformational ensemble. Specifically, we speculated that phosphorylation primarily affects LIM availability to the Z1Z2 segment by controlling the extension of the linker that connects the LIM domain and the villin headpiece. We propose that this sets the local concentration of the LIM domains near the titin N-terminus (Fig. 12A). Hence, this proposed model suggests that the association of the villin headpiece with actin could allow the LIM domains to bind to the Z1Z2 segment, with the potential to compete with or otherwise disrupt the interactions between telethonin and the Z1Z2 segment. Our docking results surprisingly indicated that phosphorylation between LIM domains does not have a significant impact on the interactions of the Z1Z2 segment, apparently in contrast to previous findings from an in vitro binding assay [3]. However, computational limitations precluded us from docking larger constructs, for which multisite phosphorylation could, in principle, disrupt its interaction with the small titin segment. It is also possible that the association of the Z1Z2 segment with ABLIM1 may also be controlled by the specific phosphorylation pattern within the ABLIM1 disordered C-terminus. The ELM database suggests that ABLIM1 has numerous potential GSK3*β* phosphorylation sites, of which several were found to be phosphorylated by mass spectrometry [3]. In addition, a study on the recruitment of ABLIM1 to cortical actin found that phosphomimetics at S214 and S431 were sufficient for ABLIM1 to co-localize with the cell membrane [7], evidencing that sites both within and outside the region containing the LIM domains have functional phenotypes. Changes in ABLIM1 phosphorylation could also impact LIM/actin interactions among other targets of the LIM domain and thus influence localization in vivo. Therefore, it is important to quantify the full range of potential phosphorylation sites in human ABLIM1 to appreciate the phosphorylation- and GSK3*β*-dependence of titin association.

### 5.3 Consequences in vivo

Based on our ABLIM1 simulation data, we aimed to relate the interfilament spacing in the sarcomere to the chains’ end-to-end distance. Our calculations revealed that the LIM domains of the full-length protein would likely be found approximately 82 Å from the C-terminal villin headpiece when phosphorylated versus 93 Å when not. Hence, the ABLIM1 LIM-Z1Z2 segment interaction would be most likely when the disordered domain spanning the N-terminal LIMs and C-terminal villin is not phosphorylated. In principle, if ABLIM1 were tethered to actin by the villin headpiece, the LIM domains could interact with the titin Z1Z2 segments on an adjacent thin filament, given interfilament distances of approximately 30 nm reported in [52]. Moreover, as the sarcomere lattice contracts under isovolumetric stretching [62], the likelihood of interaction would further increase. This mechanism is analogous to the role of phosphorylation in toggling interactions between the N-terminal extension of MyBPC and actin, where the protein is tethered to the thick filament by its C-terminal domains [63].

### 5.4 Length-Dependent Activation (LDA)

Phosphorylation’s ability to modulate interactions with the Z1Z2 segment could potentially influence myofilament mechanics. We believe that the most immediate impact would be reflected in the length-dependent activation of the sarcomere, which describes how more force is generated as the sarcomere is stretched. This phenomenon is a fundamental part of the Frank-Starling effect, where fewer equivalents of calcium are needed to achieve equivalent numbers of cross-bridges relative to when the sarcomere is at its resting length. A recognized contribution to LDA is the passive tension arising from titin, which helps restore the filaments to their equilibrium length when stretched [29]. It has been suggested that this passive tension enhances the recruitment of myosin in the ‘off state’ to the ‘on state’; the myosin ‘on state’ facilitates the formation of force-generating cross-bridges with the thin filament [29]. We postulate that some of this passive tension is due to titin binding to actin termini through dimerization of their Z1Z2 segments through telethonin, which is supported by Pinotsis *et al* [64]. In turn, factors that attenuate this anchoring are expected to decrease LDA, as has already been argued for titin treatments that disrupt passive tension [65]. Mutations in the Z1Z2 segment have also been shown to reduce its affinity for telethonin by up to 40%, which would likely impair passive tension [66] Hence, we speculate that interactions (such as the association of the LIM domain) that interfere with the binding of the Z1Z2 segment to telethonin could also have an impact on LDA.

Consistent with this idea, recent studies [9, 67] suggest that phosphorylation during heart failure decreases passive tension. In addition, GSK3*β* phosphorylation was shown to reduce ABLIM1 binding to the titin Z1Z2 segment, suggesting a tightly regulated balance between achieving an extended ABLIM1 conformation to promote titin/ABLIM1 interactions and directly modulating binding to the Z1Z2 segment, possibly through direct interactions of the LIM domain. Along these lines, telethonin deficiency leads to animals being less adaptive to stress due to reduced fractional shortening [68].

Using the Campbell myofilament contraction model, we fit its parameters to resemble the pCa50 data from Stachowski *et al*, in part by reducing the myosin off-to-on transition rate owing to reduced passive tension for the GSK3*β* KO case. Our simulation results showed a markedly reduced ΔpCa50 in the GSK3*β* knockout condition (0.05 compared to 0.26 in WT). The reduction in ΔpCa50 supports the hypothesis that decreased passive tension underlies the observed loss of LDA. Although the study by Stachowski et al. did not explicitly examine the effects of ABLIM1 phosphomimetics on LDA, evidence from MYBPC phosphomimetics and troponin I suggests that aspartic acid substitutions could reproduce the effects of phosphorylation on LDA [69].

A direct connection between ABLIM1 regulation and cardiac dysfunction remains to be established. However, in a canine model of heart failure, decreased GSK3*β* activity was reported, which was associated with a reduction in ABLIM1 phosphorylation [9]. Similarly, ABLIM1 knockdown in C2C12 myoblasts has been shown to decrease serum response factor-dependent transcription, indicating a role for ABLIM1 in sarcomeric stress signaling [70]. It should be noted that the potential interference with the binding of the Z1Z2 segment is just one of many mechanisms by which passive tension can be regulated. For example, there is a precedent for accessory proteins such as Ankrd1 and FHL2 to modulate passive tension by partitioning the I-band region of titin away from the thin filament [71–73]. Although this partitioning mechanism appears critical for these specific proteins, it has not yet been evaluated for ABLIM1. Consequently, our hypothesis that ABLIM1 may instead function by interfering with the binding of the titin Z1Z2 segment to telethonin (Tcap) remains the most consistent explanation based on available data.

### 5.5 Modulation of IDR properties

The intrinsically disordered regions within ABLIM1 are important in regulating myofilament function. One attribute is its ability to exist as an ensemble of conformations, which allows it to interact with multiple components of the myofilament, including actin and myosin, and modulate their activities in response to changing conditions, including pH and ionic strength [1]. This is because of the rugged energy landscape of ABLIM1 that enables it to sample a wide range of potential functional states. This property is particularly advantageous in the context of muscle contraction, where ABLIM1 can rapidly adapt to changing conditions to maintain optimal contractility.

PTMs provide a basis for many proteins in modulating the accessibility of binding sites and surfaces important for recognizing protein targets [74]. Phosphorylation, in particular, alters the charge distribution of ABLIM1, shifting its conformational ensemble and influencing its interactions with other proteins, as demonstrated in our study. Supporting this are data suggesting that mutations S214A and S431A in ABLIM1 have been shown to disrupt cancer migration [7], potentially by disrupting its interactions with the actin cytoskeleton. Along these lines, we previously identified [1] 92 PTM sites in the PhosphoSitePlus, database, of which the majority were phosphorylations, but we also identified acetylation and ubiquitination sites. Similar to PTMs, mutations in IDR have the potential to impact the conformational ensemble or target binding. It is not surprising that many mutations are found in disordered regions of myofilament associated protein with intrinsic disorder (MAPID)s, as reported in a recent review [1]. Tens of missense variants have been identified in ABLIM1 in GnomAD [75] (data not shown), although they remain uncharacterized; to our knowledge, a pathological variant has not been identified.

### 5.6 ABLIM1 has numerous SLIMs encoding for kinases

Our previous analysis of ABLIM1 [1] revealed a large number of phosphorylation-based post-translational modifications; many of which were in its intrinsically disordered regions, suggesting a complex regulatory mechanism. Coimmunoprecipitation experiments have identified numerous targets of ABLIM1, including the titin Z1Z2 segment [3]. To expand on these hits, we queried the ELM webserver [76] for short linear interaction motif (SLIM)s in ABLIM1 and cross-referenced them with phosphorylated fragments identified in mouse data from Stachowski et al. [3]. The ELM results indicate that ABLIM1 harbors dozens of potential SLIMs, many of which are localized to its N-terminal half, of which some are at the MSSSP segment (Table S3). In particular, the MSSSP sequence is a potential target for several proteins, including GSK3*β*, cyclin-dependent kinase (CDK), casein kinase 2, polo-like kinase 1, and phosphoinositide-dependent protein kinase 1 (e.g., mitogen-activated protein kinases). Beyond this region, GSK*β* phosphorylation motif sites are found throughout the ABLIM1 sequence. In line with these findings, knocking out GSK3*β* resulted in decreased, but not completely negated, phosphorylation of ABLIM1, suggesting that the altered activity of other kinases such as CDK could lead to further changes. At a minimum, these data are highly suggestive of ABLIM1 being under multiple modes of regulatory control by GSK3*β* and potentially other kinases.

### 5.7 Limitations

We acknowledge key limitations and assumptions that could motivate future investigations. First, the uncertainty surrounding the interaction between LIM domains and Z1Z2 segments makes it difficult to validate our predicted docking poses. Furthermore, rigid docking may obscure the actual binding mode that would result from proteins relaxing upon binding, which could lead to inaccurate predictions. Additionally, although we suggest that ABLIM1 can bind to the Z1Z2 segment in complement to previous studies that have established a link between GSK3*β* KO and decreased passive tension [3], a direct link remains to be determined. Resolving the number of myosin heads available for recruitment using methods such as small-angle x-ray scattering may help [52] in this regard. Also, other z-repeats within titin could in principle compensate for Z1Z2 segment/telethonin interactions potentially disrupted by ABLIM1 binding. However, the association between dilated cardiomyopathy and Z1Z2 segment mutations that reduce telethonin binding suggests that the titin Z domains do not necessarily compensate for Z1Z2 segment perturbation. **rev11a** Finally, our current atomistic simulations using phosphomimetic constructs may under-estimate the structural consequences of true phosphorylation. Nonetheless, our phosphomimetic structures qualitatively reproduce phosphorylation-induced conformation changes albeit with subtle differences, which is consistent with reports for other IDRs [77]. It remains possible that more complex structural behavior like aggregation may not be recapitulated using phosphomimetics [78].

## 6 Conclusions

The key results of our study reveal significant information about the role of ABLIM1 in the sarcomere. In particular, the LIM domains of ABLIM1 were found to be capable of binding titin Z1Z2 segment, and its intrinsically disordered chain properties were shown to depend on phosphorylation. Importantly, the protein likely spans the lattice spacing in the z-disk of the sarcomere in a phosphorylation-dependent manner. These findings have broad implications for our understanding of heart failure (HF). For example, the decrease in phosphorylation during HF [9, 67] suggests that post-translational modification (PTM) control of LIM-containing proteins may play a critical role in disease progression. Together with the observed decrease in length-dependent activation in HF versus wild-type conditions from Stachowski et al., this highlights the importance of phosphorylation in tuning the functional contribution of ABLIM1 to cardiac contractility.

## 7 Acknowledgements

The research reported in this publication was supported by the Maximizing Investigators’ Research Award (MIRA) (R35) from the National Institute of General Medical Sciences (NIGMS) of the National Institutes of Health (NIH) under grant number GM148284. This work used PSC Bridges from the Advanced Cyberinfrastructure Coordination Ecosystem: Services & Support (ACCESS) program, which is supported by National Science Foundation grants #2138259, #2138286, #2138307, #2137603, and #2138296. We are grateful for Stefano Sala’s careful reading of the manuscript.

## Supplementary Information

### S1 Supplement

#### S1.1 Supplemental Tables

**Table S1:** Phosphorylation sites from [3] (mouse)

| Site | Fold change | Modeled |
| --- | --- | --- |
| 215/216 | --- | S76 in isolated LIM |
| 354 | 0.516 (p=0.042) | Not in human |
| 494 | 0.627 (p=0.043) | In human (450, 452) |
| 496 | 0.630 (p=0.045) |  |

**Table S2:** Summary of parameters and distances. used for Flory polymer model.

| Description | Value |
| --- | --- |
| b | 3.4 for a.a [79] |
| $\nu_{unf}$ | 0.58 [23] |
| $\nu_{fld}$ | 0.34 [23] |
| $R_g$, ProTα | $\approx 3.0$ [nm] [23] |
| actin/actin separation | 30.4 [nm] [52] |
| $N_{1-97}$ | 97 |
| $N_{343-473}$ | 131 |
| $N_{343-710}$ | 368 |
| $\langle R_e \rangle$ wild-type, 343-473 | 5.3 [nm] Fig. 9 |
| Flory exponent, $\nu$, wild-type, 343-473 | 0.68 Fig. 11 |
| $\langle R_e \rangle$ S450D/S452D, 343-473, | 4.8 [nm] Fig. 9 |
| Flory exponent, $\nu$, S450D/S452D, 343-473 | 0.65 Fig. 11 |
| Flory prefactor, $R_0$, 343-473 | 0.60 [nm] |

**Table S3:**
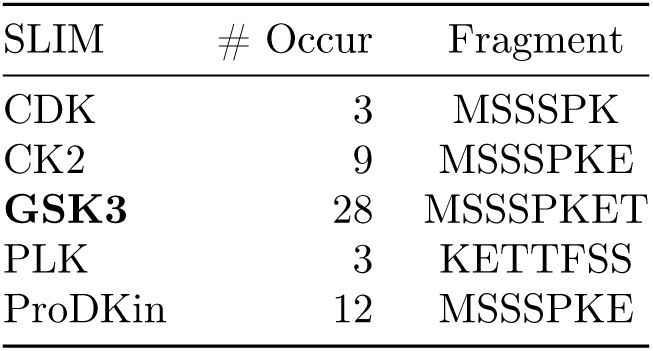
Table of short linear interaction motifs serving as kinase targets, as suggested in the ELM database [80] for human ABLIM1. # occur refers to the number of predicted motifs in the sequence, inclusive of the MSSSP sequence.

| SLIM | # Occur | Fragment |
| --- | --- | --- |
| CDK | 3 | MSSSPK |
| CK2 | 9 | MSSSPKE |
| **GSK3** | 28 | MSSSPKET |
| PLK | 3 | KETTFSS |
| ProDKin | 12 | MSSSPKE |

**Table S4:**
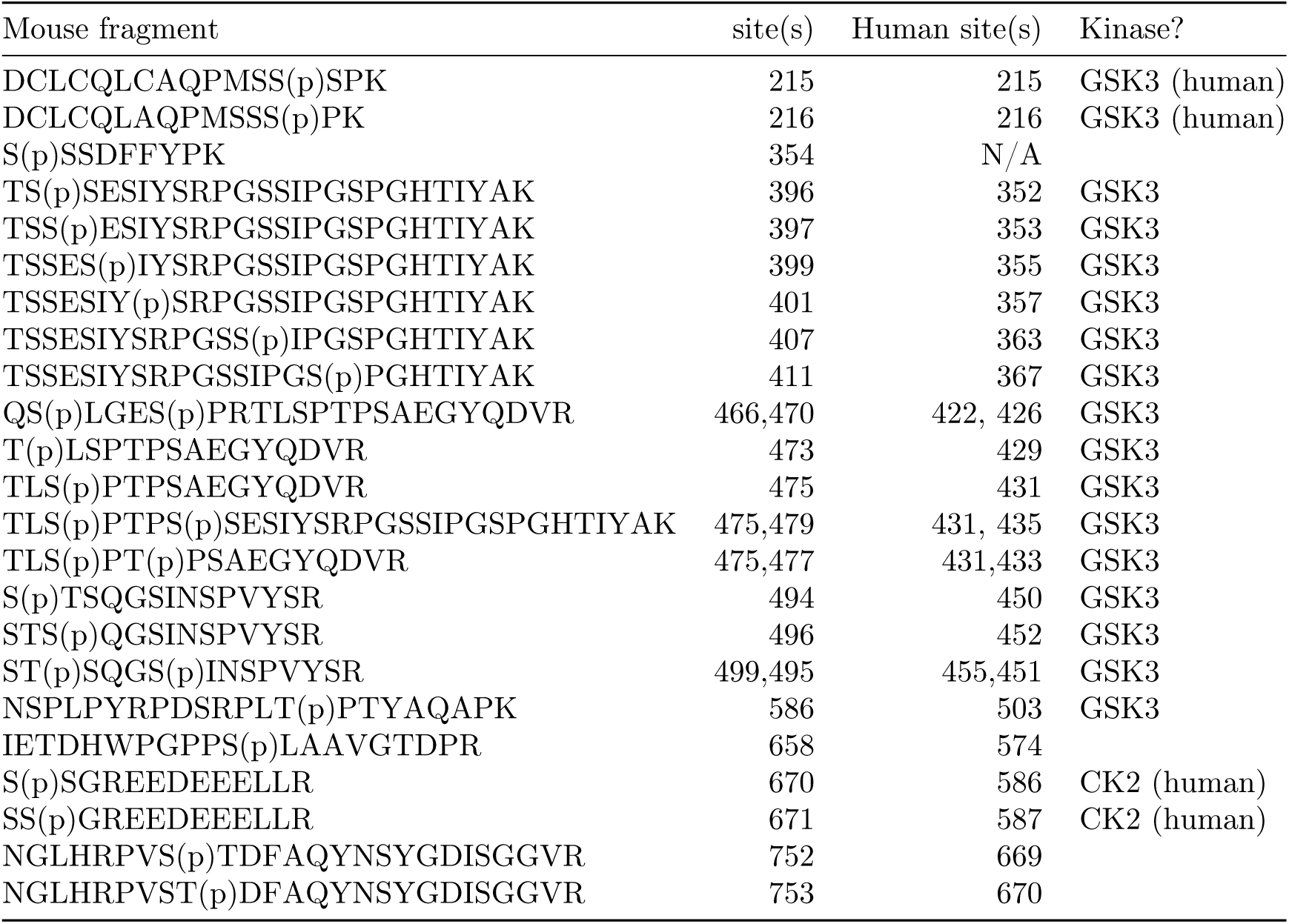
Tabulation of phosphorylated fragments identified in [3], compared to the homologous positions in human ABLIM1. Sites annotated as GSK3B phosphorylation sites by ELM are marked as such.

**Table S5:**
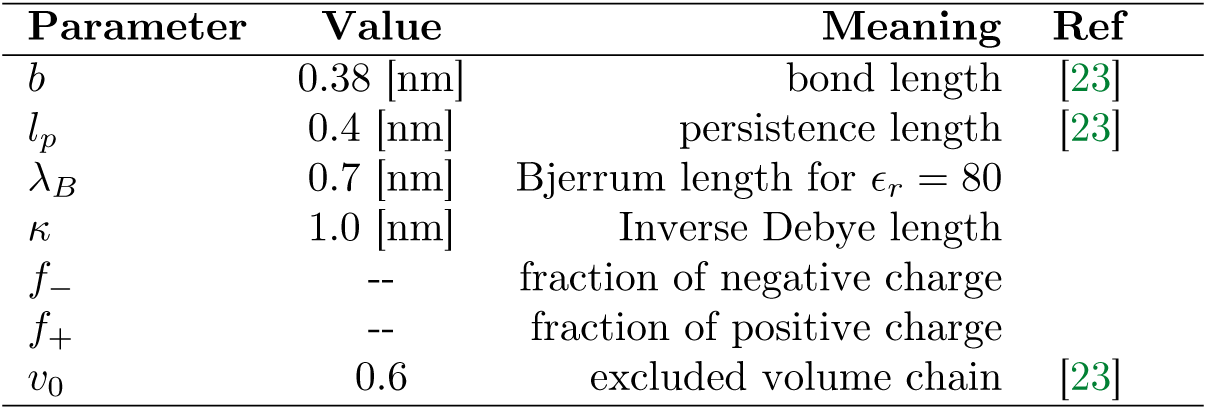
Parameters used in the polymer physics model.

**Table S6:** Data table for sites, sequences, and values of *f_p_* and *f_n_*.

| Case | Sites | Sequence | $f_p$ | $f_n$ | rep/attr |
| --- | --- | --- | --- | --- | --- |
| ABLM1 | 210-222 | AQPMSSSPKETTF | 0.07 | 0.07 | neutral |
| ABLM1 | 210-222 | AQPMdSSPKETTF | 0.07 | 0.14 | repulsive |
| MYBPC3 | c0c1 | RSAFRRTSLAGAGRRTSDSHED | 0.23 | 0.14 | repulsive |
| MYBPC3 | c0c1 | RDAFRRTSLAGAGRRTSDSHED | 0.23 | 0.18 | less repulsive |

#### S1.2 Supplemental Methods

##### S1.2.1 Parameterization of phosphorylated serine residues in C-terminus of ABLIM1

Phosphomimetics are an imperfect model of phosphorylation. For instance, the S129D phosphomimetic failed to reproduce the structural changes and aggregation observed for phosphorylated *α*-synuclein [78]; nevertheless, structural insights derived from phosphomimetic mutants often align with results from *in vitro* phosphorylation studies [43] and are commonly used to investigate phosphorylation-induced structural and functional alterations in proteins [81, 82]. The pSerine force field parameters were adapted from Amber ff19SB [41] for atomistic simulation and from Pluhackova *et al* ‘s work [42] for coarse-grained simulations. For producing the coarse-grained model, the atomistic structure of ABLIM1 was first mutated to S450K and S452K to provide a reference amino acid at these sites that Martini3 would convert into a bead with two side chains, the same number as phosphorylated serine residues. The same martinize.py script and DSSP program were used as in the original CG MD simulations 3.5, and the resulting pS450 and pS452 residues of the .itp file were modified (see Table S7 and Table S8) to reflect previous generation of Martini phosphoserine beads in the literature [42] and modified for our Martini3 algorithm instead of Martini2.2. After generating the pS450_pS452 CG structure, simulation preparation and production were performed exactly as the previous CG MD simulations were and resulted in seven 2000ns runs for analysis.

**Table S7:** Bead labelling and data for phosphoserine residues in Martini3 coarse-grained simulations.

| Residue | Bead name | Bead type | Charge |
| --- | --- | --- | --- |
| SP2 | P2 | BB | 0 |
| (phosphoserine) | SP1 | SC1 | 0 |
|  | D | PO4 | -2 |

**Table S8:**
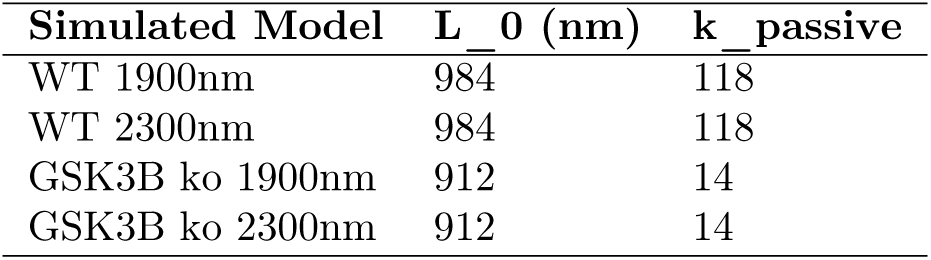
Bonded parameters for phosphoserine residues in Martini3 coarse-grained simulations.

**Table S9:**
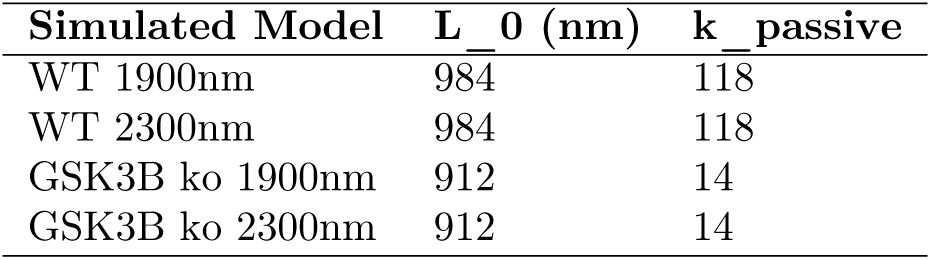
Length-Dependent Activation parameters for MATLAB simulation of stress impacts on calcium binding affinity. The parameters that were consistent for all simulation models can be found in the following table.

**Table S10:** Parameters that were changed from the default tension-pCa demo values in the model file for MATMyoSim.

| Parameter | k<sub>force</sub> | k<sub>off</sub> | k<sub>1</sub> | k<sub>2</sub> | k<sub>3</sub> | k<sub>4_0</sub> | k<sub>4_1</sub> |
| --- | --- | --- | --- | --- | --- | --- | --- |
| value | 6.17 | 50 | 1.74e-4 | 200 | 7.11 | 1.69 | 0.1 |

| Parameter | k<sub>cb</sub> | x<sub>ps</sub> | k<sub>on</sub> | k<sub>off</sub> | k<sub>coop</sub> | k<sub>linear</sub> |
| --- | --- | --- | --- | --- | --- | --- |
| value | 0.001 | 1.4 | 2.08e7 | 50 | 5.70 | 14 |

#### S1.3 Supplemental Figures

#### S1.4 Supplemental Results

##### S1.4.1 Disorder

We additionally used the globprot web interface [31] to predict intrinsically-disordered regions for ABLIM1 (ACCESSION code O14639), based on the Russell-Linding statistic [31]. The predictions for the folded LIM domains and villin head piece showing in Fig. S4 are consistent with IUPRED 2 A predictions in the main text. The disordered regions are more prevalent in the sequence spanning 336 to 743, consistent with our other results.

**Figure S1:**
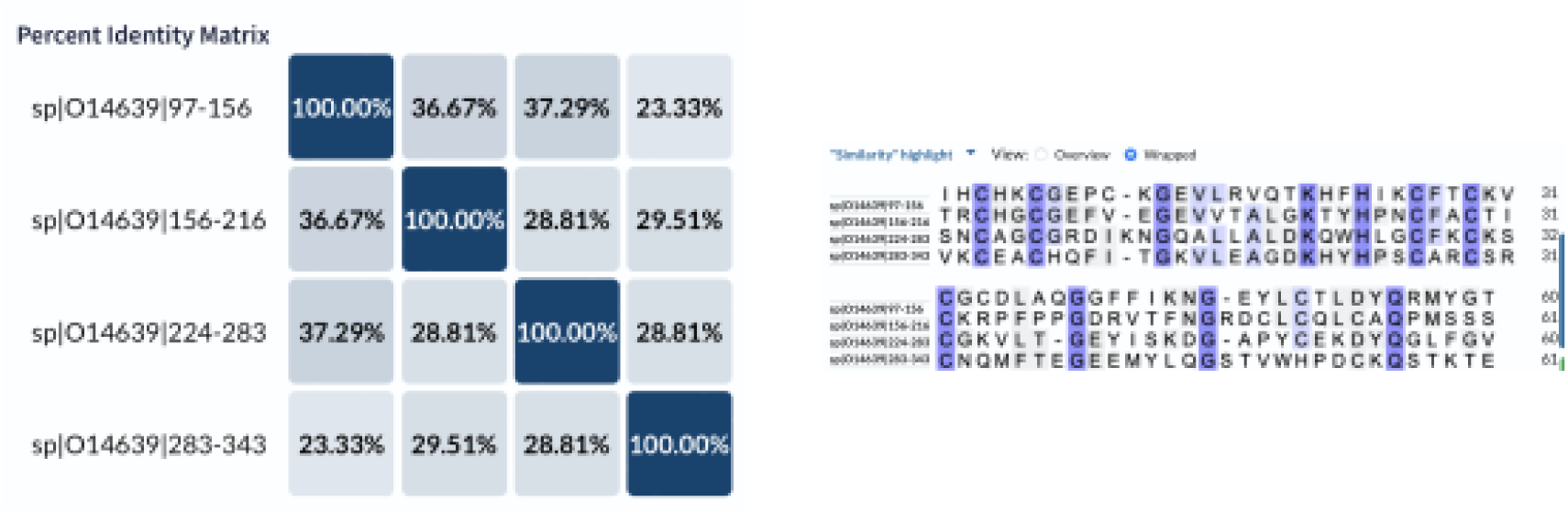
Conservation between LIM domains in ABLIM 1.

##### S1.4.2 Atomistic MD of the four LIM domain construct with and without phosphorylation at S216

In order to explore the effects of phosphorylation at S216 on the conformations of the folded LIM domains, we performed atomistic MD simulations on the wild-type (WT) system, comprising residues 97-343 of ABLLIM1, and on the phosphorylated system, where S216 was mutated to D216 to mimic the negative charges introduced by phosphorylation (Fig. S6A). MD simulations were carried out using the Amber20 package [83], with the protein described by the ff19SB force field [84] and solvated in a 10 Å margin OPC water box [85], containing 0.15 M KCl salts. The simulation protocol included energy minimization, heating, equilibration, and a 300 ns production run, with the detailed protocol reported in our previous study [86]. During the MD simulations, the coordination between the Zn^2+^ ions and their coordinating residues was maintained using harmonic restraints with a force constant of 20 kcal/mol/Å^2^. Since the simulations were initiated from the AlphaFold-predicted structure, we observed that during the 300 ns production run, both the WT and S216D systems underwent significant conformational changes and became stable by the end of the simulation, with RMSDs of approximately 20 Å compared to the initial structure. To compare the conformational ensembles between WT and S216D, we calculated the radius of gyration (R*_g_*) for both systems (Fig. S6B). We found that the S216D system experienced significant expansion and collapse in its conformation, as evidenced by an increasing and then decreasing R*_g_* curve, whereas the WT system showed a relatively stable R*_g_*. As a result, the R*_g_* distribution for S216D spanned a wider range (25–50 Å) compared to the WT system (25–40 Å). These findings from the atomistic MD simulations agree with the observations made in our coarse-grained (CG) simulations. Phosphorylation at S216, located in the linker region that connects the LIM2 and LIM3 domains, can indeed affect the conformation of the entire folded region of ABLIM1, promoting a more open conformation.

##### S1.4.3 Fractional charge and self-repulation

The Flory ideal chain model predicts that the radius of gyration scales as a function of *ν*[23]

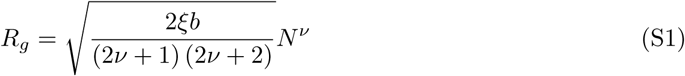

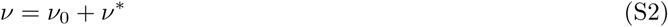

Deviations from ideal Flory scaling of polymer properties have in part been attributed to imbalances in the fraction of positively versus negatively charged amino acids [23].

**Figure S2:**
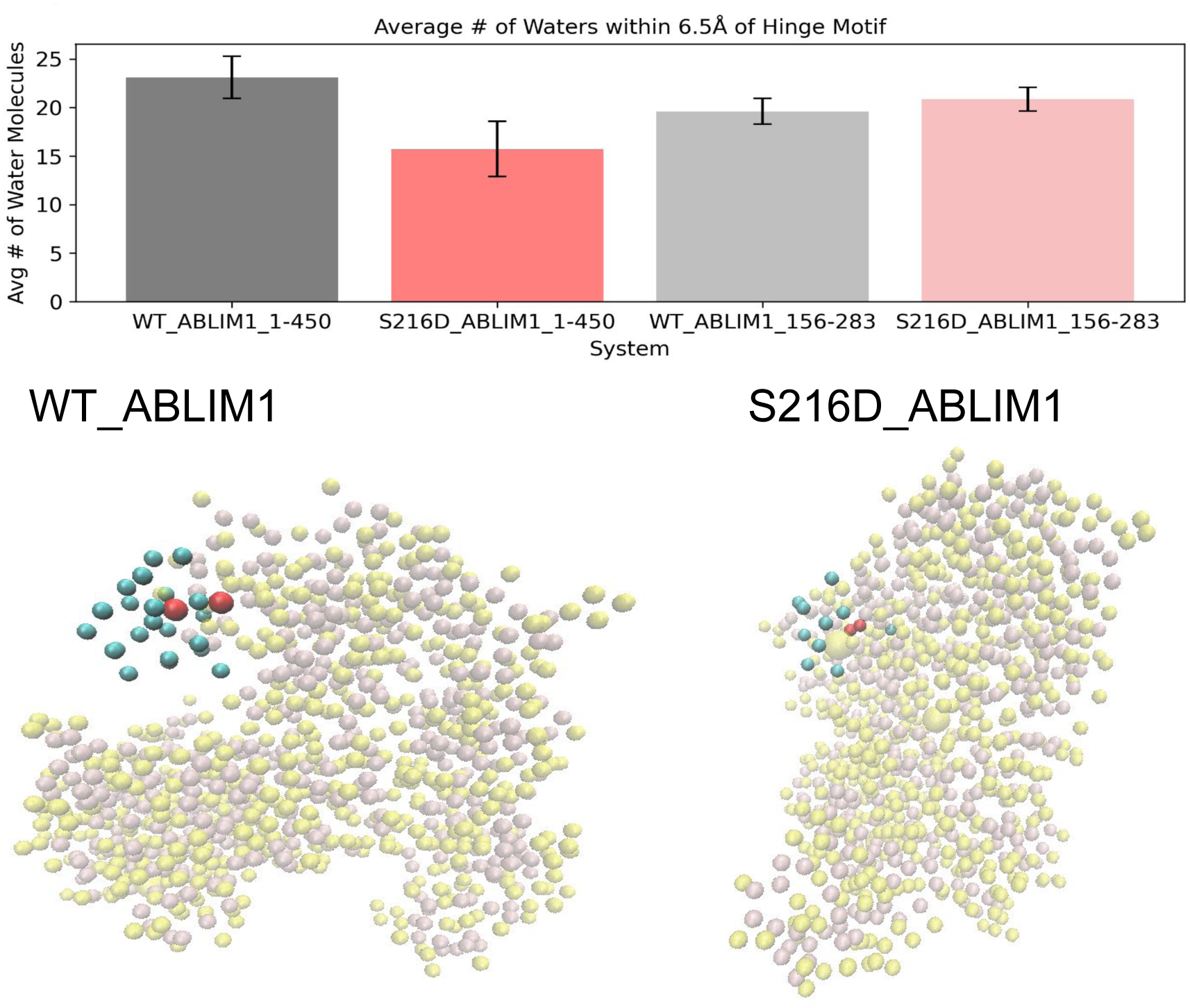
Average number of water molecules within 6.5Åof the hinge motif (Res 212-220) in coarse-grained ABLIM1 models

Attractive versus repulsive corrections to scaling coefficient *ν* can be estimated using

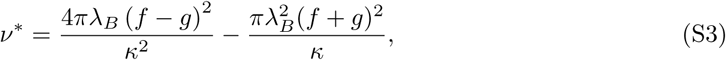

for which we define its parameters in Table S5. According to Hofmann supp, *ν*^∗^ *>* 0 refers to net repulsion (unfolding that increases the effective volume), while *ν <* 0 corresponds to net attraction (folding).

As an example for the effects of fractional charge density, phosphorylation of ABLIM1 in the LIM2-LIM3 loop increases the fractional negative charges, which potentially extends the disordered region (see Table S6). In contrast, the C0C1 region in MYBPC3 has a higher fractional density of positive residues relative to negative, which we expect to extend the disordered region. Its phosphorylation would decrease the difference in charge density and potentially contract the chain. Indeed, phosphorylation has been shown to reduce MYBPC’s interaction with the thin filament, suggesting that the structure contracts [87]. This is supported by findings from the Warshaw lab that phosphorylated MYBPC3 has a shorter contour length than the wild-type [88].

**Figure S3:**
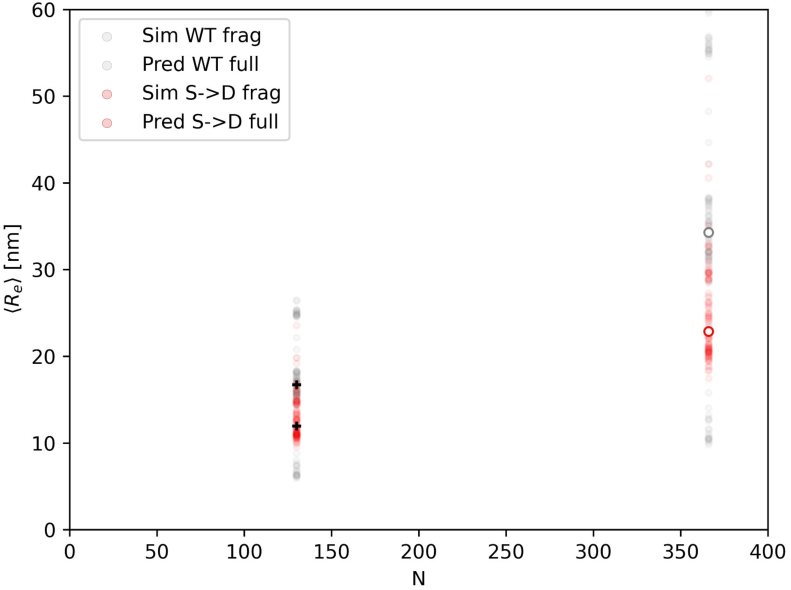
Predictions of ⟨*r_e_*⟩ with respect to number of amino acids (N), based on fitting to fragment simulation data using Eq. 1.

**Figure S4:**
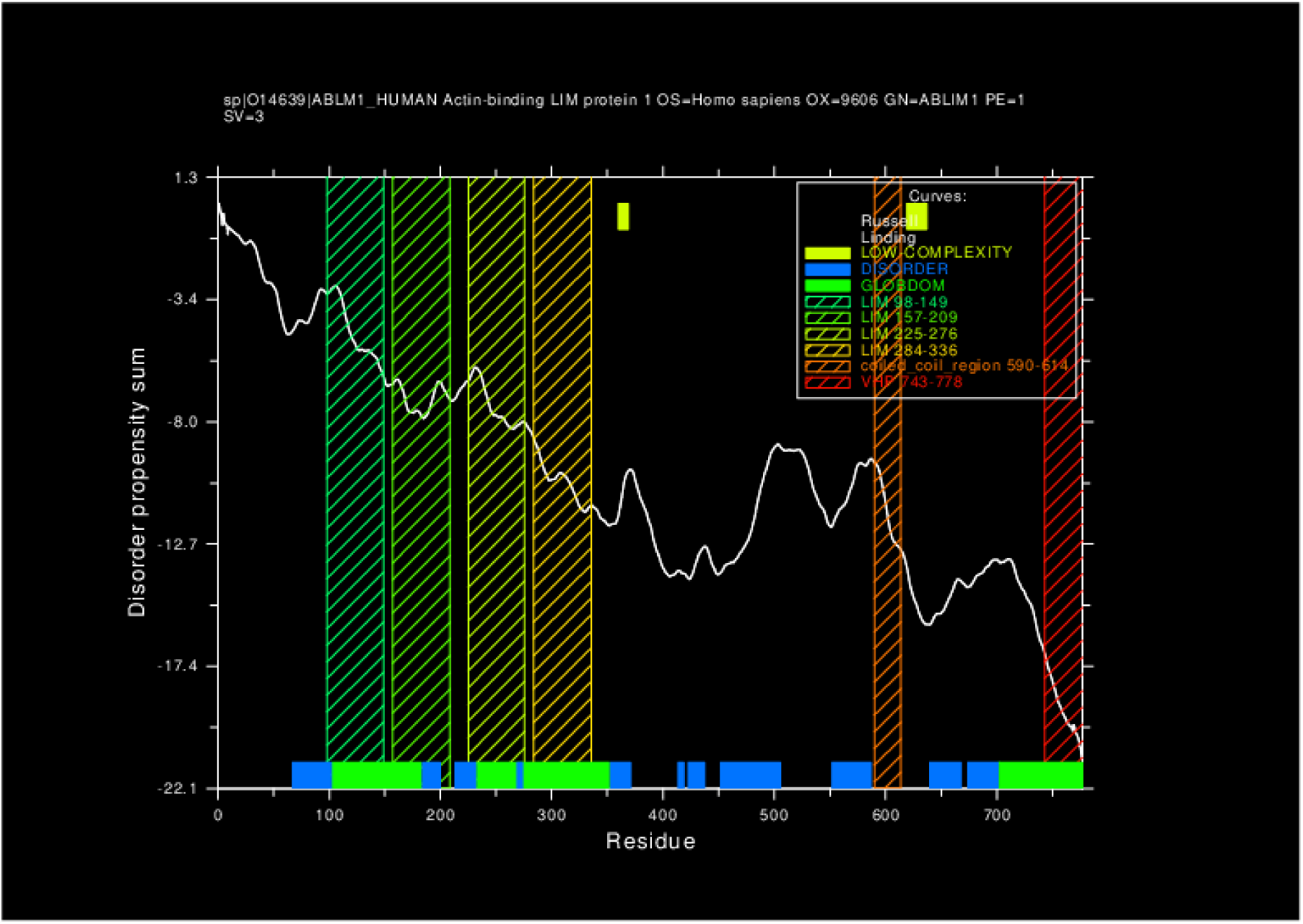
Predictions of disorder in ABLIM1 using the globprot interface [31].

**Figure S5:**
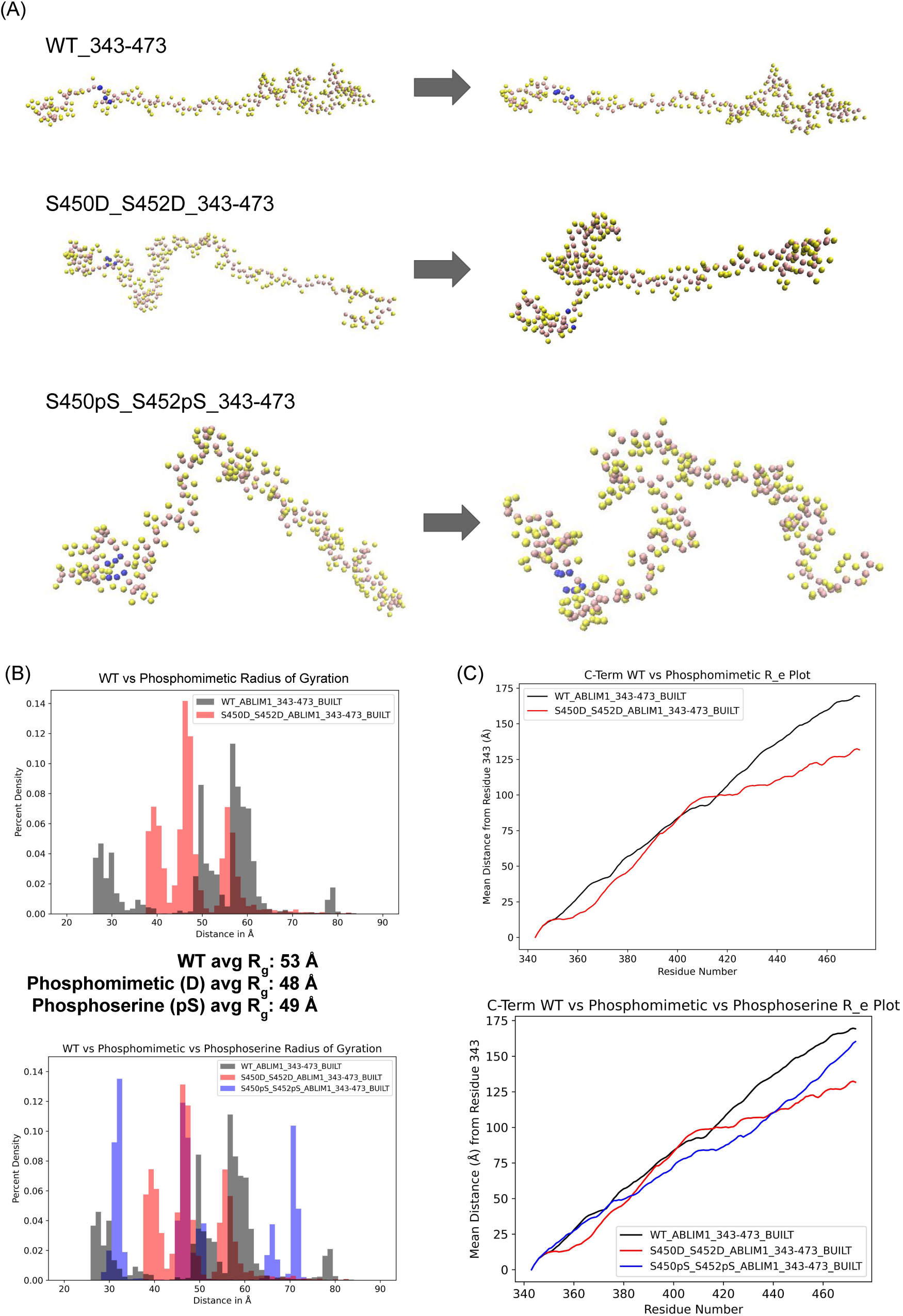
Comparison of the double phosphoserine protein with the existing WT and phosphomimetic constructs. A) Representative structures at beginning (∼300ns) and end (∼1500ns) of production simulation. B) Radius of gyration in all systems and their computed average R*_g_* across all simulations. The top figure is presented above in the paper while the bottom has added the phosphoserine simulation runs for comparison. C) Average end-to-end distance across all C-terminus simulations.

**Figure S6:**
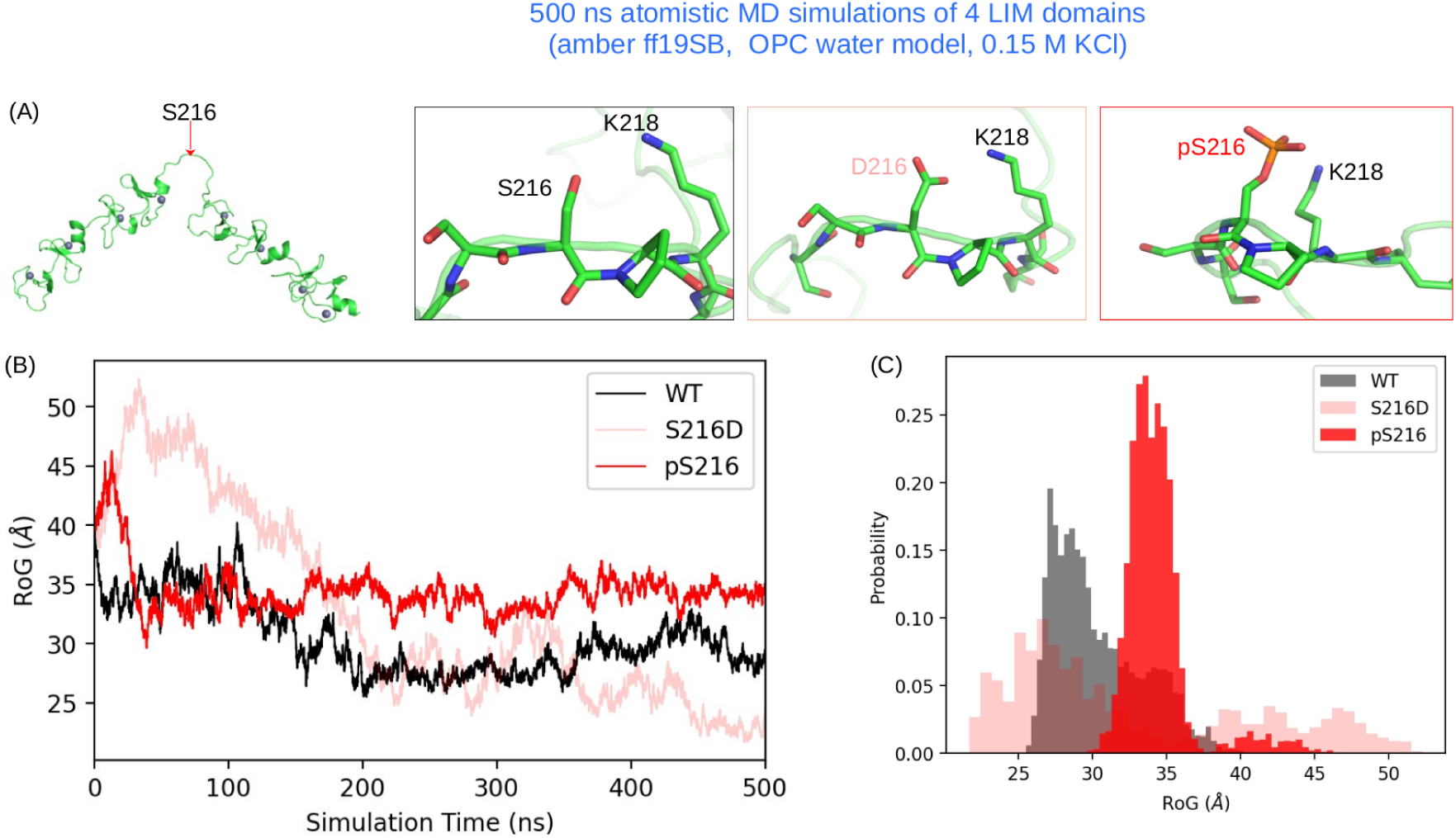
Atomistic MD runs of WT and phosphorylated ABLIM1 four LIM domain constructs. We performed MD simulations on the phosphomimetic S216D and the actual phosophorylated S216 (pS216) constructs to explore if using phosphomimetic S216D can reflect the impacts of phosphorylation on the conformational property of LIM domain construct. A) The ABLIM1 structure of residues 97-343 predicted by AlphaFold was used as starting structure of MD simulations. The location of S216 in between LIM2 and LIM3 domains is indicated by a red arrow. The Zn^2+^ cations are depicted as gray spheres. B) Time-dependent RoG (R*_g_*) and (C) the distributions of R*_g_* are plotted.

**Figure S7:**
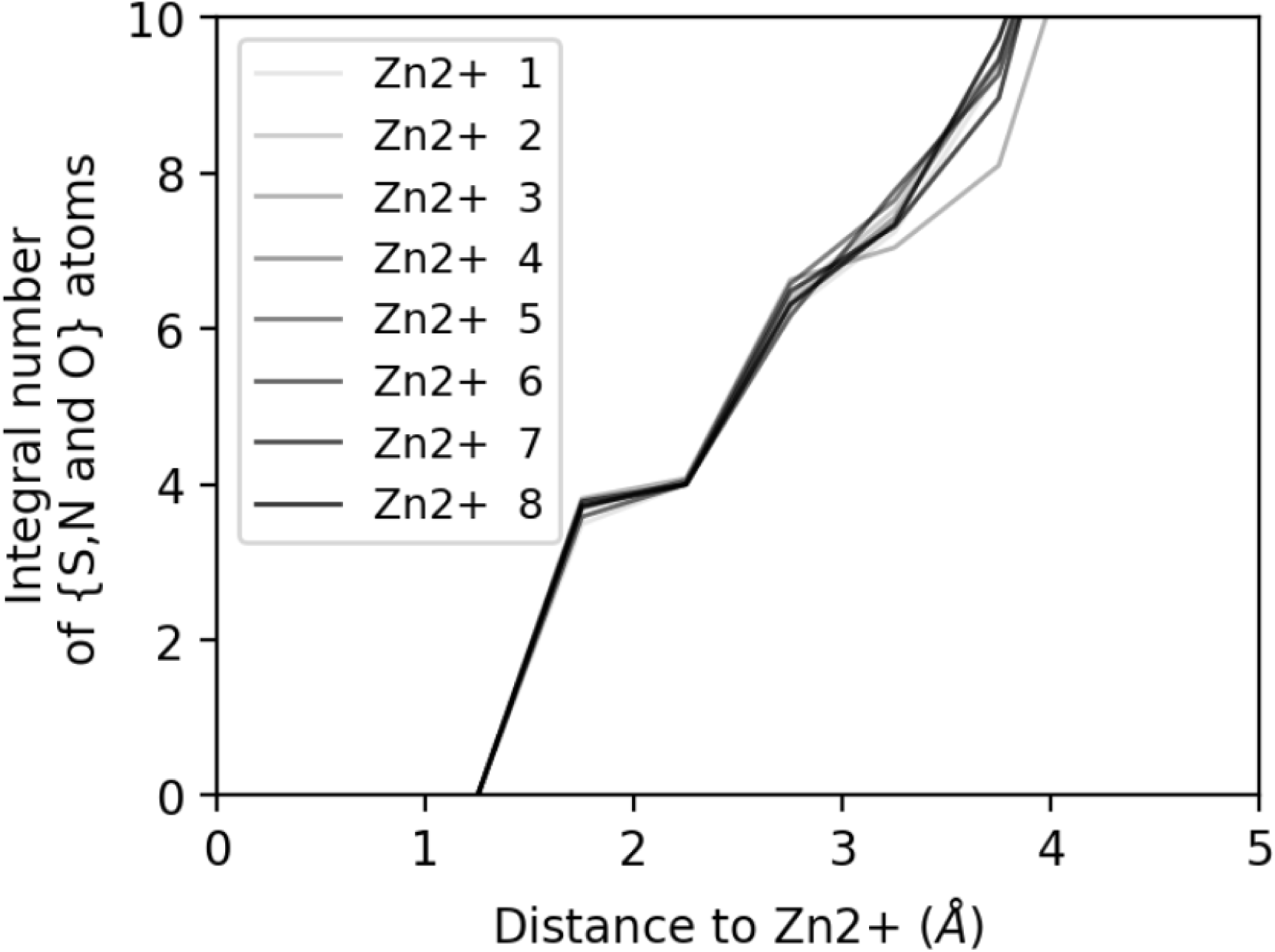
Coordination of Zn^2+^ calculated from 500 ns atomistic MD of the WT four LIM domain construct. We showed that, by introducing appropriate restraints as we discussed in the Method section, all eight Zn^2+^ ions were well-coordinated by four atoms each, reaching the theoretical coordination number of 4 for Zn^2+^.

**Figure S8:**
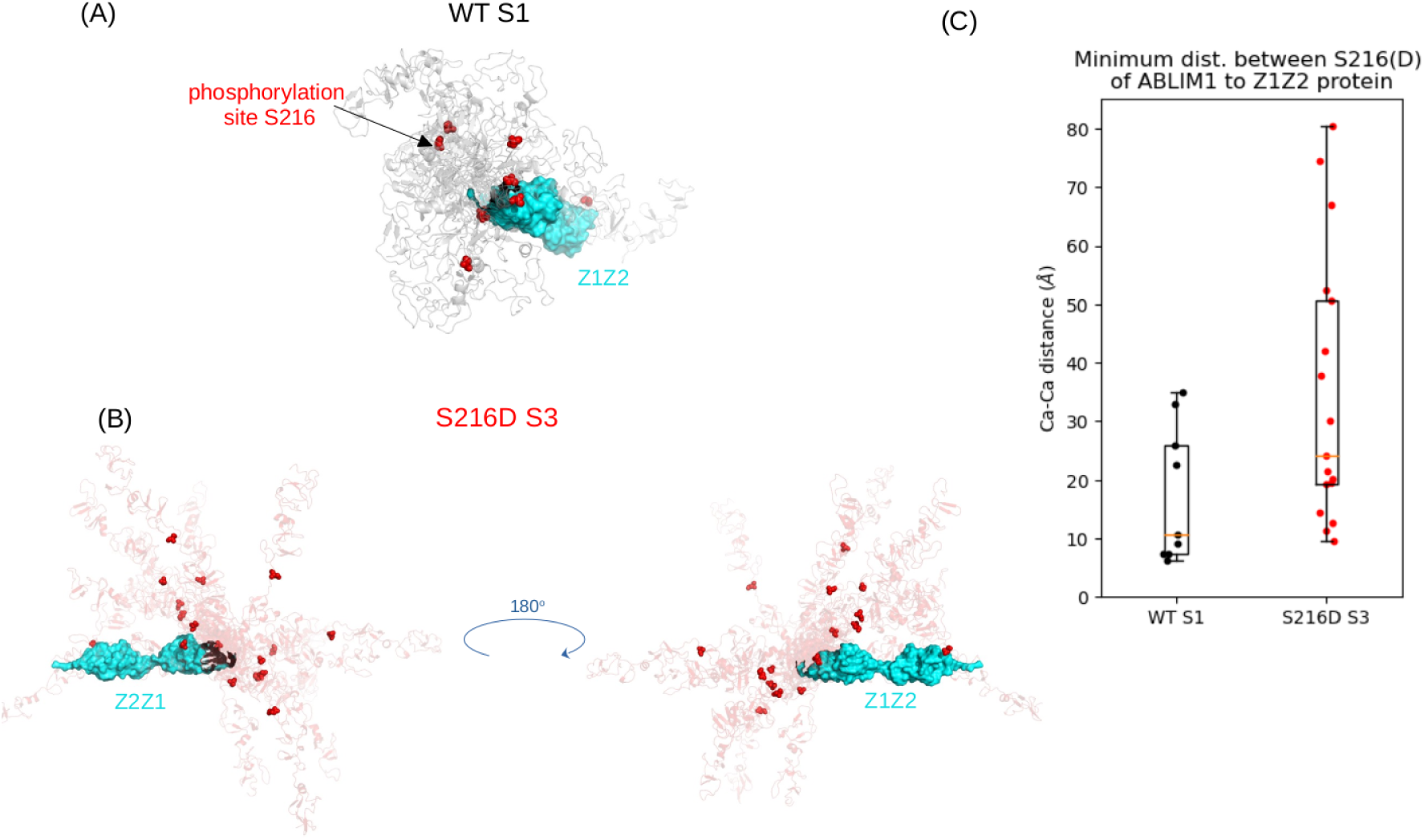
(A-B) Location of the phosphorylation site S216 relative to ABLIM1-Z1Z2 binding interface as revealed by rigid protein docking via Cluspro. (C) The minimum distance between C*α* atom of S216(D) to C*α* atoms in Z1Z2.

